# Attenuation of chronic antiviral T cell responses through constitutive COX2-dependent prostanoid synthesis by lymph node fibroblasts

**DOI:** 10.1101/457127

**Authors:** Karin Schaeuble, Hélène Cannelle, Stéphanie Favre, Hsin-Ying Huang, Susanne G. Oberle, Dietmar Zehn, Sanjiv A. Luther

**Affiliations:** Center for Immunity and Infection Lausanne, Department of Biochemistry, University of Lausanne, Epalinges, Switzerland; Swiss Vaccine Research Institute, Epalinges, Switzerland; Division of Immunology and Allergy, Department of Medicine, Lausanne University Hospital, Lausanne, Switzerland; Division of Animal Physiology and Immunology, School of Life Sciences Weihenstephan, Technical University of Munich, Freising, Germany

**Keywords:** lymph node, fibroblastic reticular cells, cyclooxygenase, prostaglandin E_2_, chronic T cell response, T cell suppression

## Abstract

Fibroblastic reticular cells (FRC) of lymphoid T zones actively promote T cell trafficking, homeostasis and expansion, but can also attenuate excessive T cell responses via inducible nitric oxide and constitutive prostanoid release. It has remained unclear under which conditions these FRC-derived mediators can dampen T cell responses and whether this occurs *in vivo*. Here we confirm that murine lymph node FRC produce prostaglandin E_2_ (PGE_2_) in a cyclooxygenase-2 (COX2)-dependent and inflammation-independent fashion. We show that this COX2/PGE_2_ pathway is active during both strong and weak T cell responses, in contrast to nitric oxide which only comes into play during strong T cell responses. In chronic infections *in vivo,* PGE_2_-receptor signaling in virus-specific CD8 T cells was shown by others to suppress T cell survival and function. Using CCL19cre x COX2^flox/flox^ mice we now identify CCL19cre^+^ FRC as the critical source of this COX2-dependent suppressive factor, suggesting PGE_2_-expressing FRC within lymphoid tissues are an interesting therapeutic target to improve T cell mediated pathogen control during chronic infection.

## Introduction

Lymph nodes (LN) are secondary lymphoid organs (SLO) specialized in filtering lymph fluid and initiating T and B cell responses to foreign antigens. Typically, antigen-specific T cells are selected to expand and differentiate into effector cells by dendritic cells (DC) that present processed antigen in the context of MHC and costimulatory signals. However, T cells may also be tolerized if self-antigens are presented in a non-immunogenic context, typically by immature DC. All of these processes take place within the T cell rich compartment of SLO which are organized by FRC that are the most prevalent non-hematopoietic cell type in this zone and play active immune regulatory roles (1, 2).

Initially, FRC have mainly been described to enhance adaptive immunity in several ways. FRC constitutively produce the chemokines CCL19 and CCL21 responsible for attracting and retaining dendritic cells and naïve T cells in that compartment and thereby facilitating their physical interaction (3, 4). These encounters are further enhanced by the dense reticular network formed by FRC, allowing DC adhesion and physical guidance for migrating T cells during their continuous search for antigen (5, 6). FRC play yet another important role in promoting adaptive immunity by producing the T cell survival factor interleukin-7 (IL-7) (3).

More recently, evidence has accumulated for negative regulatory roles of FRC in T cell responses (1, 2). FRC were shown to express self-antigens in the context of MHCI (7) or to acquire antigens and MHCII molecules from neighboring DC (8) with evidence suggesting induction of peripheral T cell tolerance. However, FRC can also inhibit T cell responses as bystander cells, presumably without need of antigen presentation. Ex vivo FRC were demonstrated to express inducible nitric oxide synthase (iNOS) upon detection of IFNγ and TNFα derived from T cells shortly after priming, leading to a nitric oxide (NO)-dependent attenuation of T cell proliferation (9-11). This mechanism was shown to be limited to the early phase of T cell priming, leading to a reduced expansion of T cells that were nevertheless functional. While strong *in vivo* evidence for a critical role of iNOS or self-antigen in FRC dampening T cell responses is still scarce, these findings suggest that FRC may protect tissues against damage caused by very strong and potentially pathogenic T cell responses.

During our initial studies with FRC lines, we observed that part of the inhibitory activity was dependent on cyclooxygenase (COX) enzymes, as it could be blocked by the pharmacological inhibitor indomethacin. Interestingly, COX2 expression by our LN FRC line was found to be constitutive, in contrast to the cytokine-induced iNOS expression (11). This initial finding suggested that FRC may have more than one pathway capable of dampening T cell responses, but the relative roles of COX versus iNOS in FRC remained unclear, as did the precise context, mechanism and *in vivo* role of COX2 specifically in FRC.

COX enzymes are the rate-limiting enzymes in the pathway converting the lipid arachidonic acid into various prostanoids (COX-dependent lipids), namely prostaglandins and thromboxane A2. There are two genes coding for the two COX enzymes that differ in their expression pattern but perform the same function. Typically, COX-1 is constitutively expressed in several cell types to support homeostatic processes, while COX2 is upregulated transiently during inflammatory reactions in injured tissue cells or infiltrating myeloid cells. Among the COX-dependent lipids, prostaglandin E_2_ (PGE_2_) was demonstrated to have powerful pro-inflammatory activities, responsible for the development of many hallmarks of inflammation, including fever, pain and swelling (12-14). Many nonsteroidal anti-inflammatory drugs (NSAID), including aspirin, ibuprofen, indomethacin and paracetamol, target this function of the two COX enzymes. They are among the most widely used drugs and have become a standard treatment for many inflammatory reactions. However, several reports have highlighted the dual function of COX-dependent prostanoids in inflammation (12-16). While during acute inflammation PGE_2_ enhances innate immunity, in part by inducing local vasodilation and attraction of neutrophils and macrophages, PGE_2_ can be very prevalent during chronic inflammation, such as cancer and persisting viral infection, when it is often observed to be anti-inflammatory and suppressing type I immunity (17-19). In addition, PGE_2_- induced effects appear to be strongly dose- and context-dependent, with T cell suppression seen for example mainly with high PGE_2_ concentrations (14, 20). These findings emphasize the importance of a better understanding of the cells expressing COX-dependent prostanoids, within SLO and target tissues, and their influence on immune response regulation *in vivo*, eventually also to improve the knowledge on mechanisms of action of the so widely used COX-inhibiting drugs.

In this study, we show that LN FRC constitutively express high levels of COX2 and its product PGE_2_, thereby dampening T cell expansion *in vitro*. This property of FRC is independent of the strength of the inflammatory stimulus, in contrast to the second negative regulator, nitric oxide. Importantly, using mice selectively lacking COX2 expression in FRC we show that this cell type and pathway is critical to attenuate T cell responses during persisting viral infection. These findings suggest that the EP2/4-dependent mechanism of T cell suppression previously observed by Kaech and colleagues during chronic LCMV infection (18) is mediated by COX2^+^ LN FRC pointing to a previously unappreciated role for FRC as negative regulators of chronic immune responses.

## Results

### Lymph node FRC dampen CD8+ and CD4+ T cell responses *in vitro* via constitutive production of COX2-dependent prostanoids

We have previously described the COX-dependent attenuation of T cell expansion in a co-culture system where T cells were activated by antigen-loaded DCs in presence of FRC (11). To exclude a role for DC as source of COX-dependent prostanoids and to study more specifically the role of COX2 in FRC, we activated naïve murine T cells using αCD3/28- coated beads in presence or absence of the FRC line called pLN2. In presence of FRC CD8+ and CD4+ T cell proliferation was strongly attenuated, as assessed by analyzing the percentage and absolute numbers of proliferating T cells, and this inhibitory effect could be reversed to a similar extent by adding either the iNOS inhibitor 1400W or the COX1/2-inhibitor indomethacin (Fig. 1A). The inhibitors enhanced the CD4+ and CD8+ T cell expansion around 3-5fold with the response reaching 35-70% of the T cell response observed in absence of FRC. While FRC typically attenuated 85-95% of the T cell response in this co-culture assay, these pharmacological inhibitors reduced this attenuating effect by a factor of 2-3 fold, allowing many T cells to undergo several rounds of cell division and to express more of the activation markers CD44 and CD25 (Fig. 1A, B, S1A). Interestingly, among the proliferating T cells there were only small differences observed regarding CD44 and CD25 expression in the various settings (Fig. 1B, S1A) suggesting FRC-expressed iNOS and COX2 reduce the number of T cells getting primed, with less effects on T cells once they have successfully entered cell cycle.

**Figure 1:**
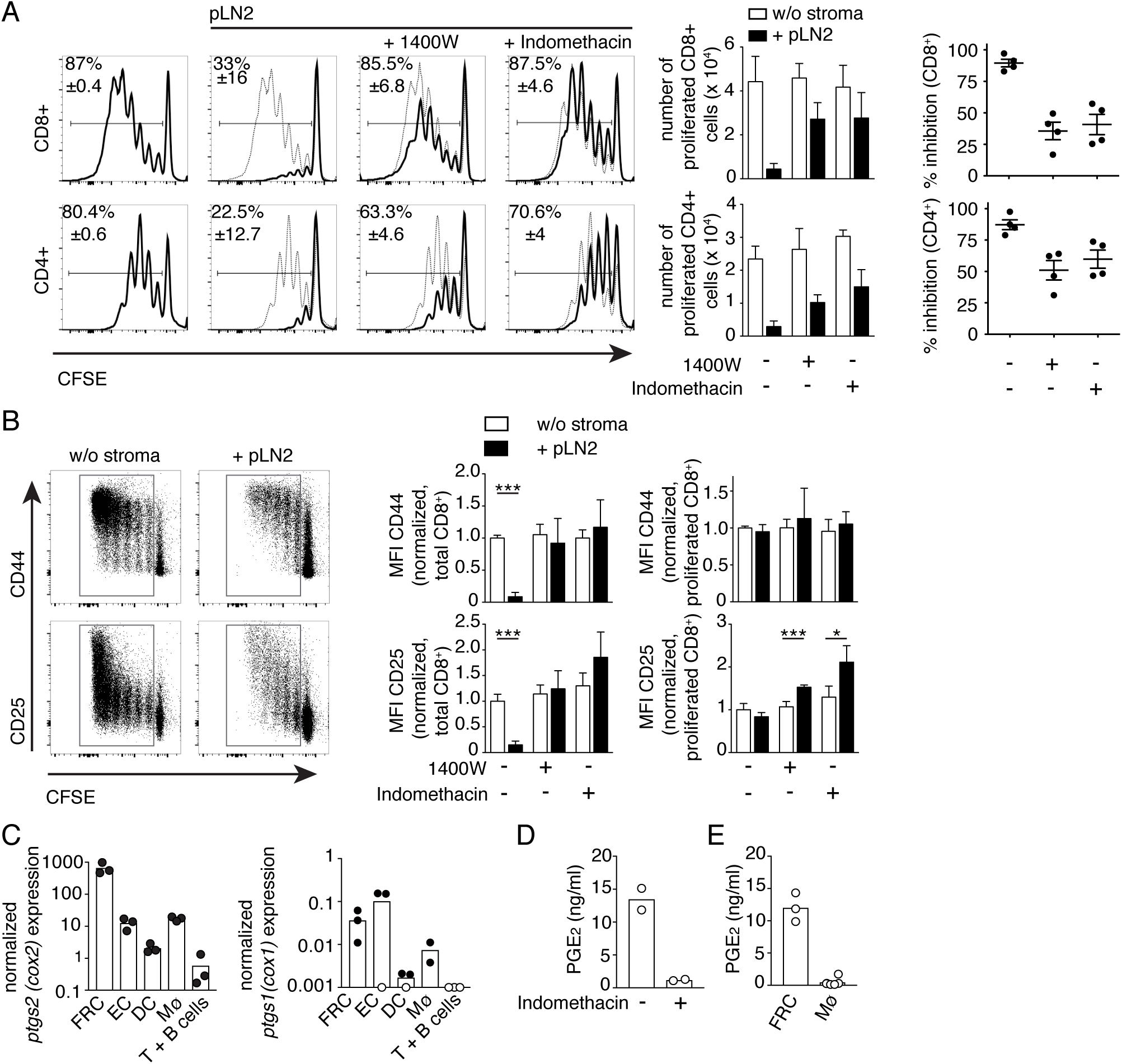
Naïve LN FRC dampen CD8 and CD4 T cell activation and proliferation by constitutive COX2 and PGE_2_ expression. (A-B) CFSE labeled T cells were activated non-specifically with αCD3/28 DynaBeads and cultured for 3d in the absence or presence of pLN2 (pLN FRC cell line) and 1400W (3µM) or indomethacin (10µM), respectively, followed by flow cytometric analysis. (A) On the left side histograms of CD8+ and CD4+ T cells showing CFSE dilution as readout of T cell proliferation in presence (thick line) or absence (thin line) of FRC. Bar graphs in the middle indicate the number of proliferated CD8+ and CD4+ T cells in presence (black bars) or absence (white bars) of FRC. Scatter plots on the right side represent the percentage inhibition of T cell proliferation mediated by FRC, calculated based on the number of T cells 3d after co-culture. (B) Dot plots showing CD44 and CD25 expression of CFSE labeled CD8+ T cells after 3 days of co-culture. Bar graphs depicting the median fluorescence intensity (MFI) of CD44 and CD25 on total or proliferated CD8+ T cells. (C) Quantitative RT-PCR analysis for *ptgs1/2* transcript levels in the indicated sorted cell types from pLN of naive wt mice (n=2-3, each sample represents a pool of 2-3 mice). Transcript levels below the detection limit or nonspecific transcripts are indicated as white circles on the x-axis. (D-E) PGE_2_ levels in the supernatant after overnight culture as determined by ELISA. (D) pLN2 cells in the presence or absence of indomethacin (10µM). (E) FRC (CD45- CD31- Podoplanin+) and CD11b+ macrophages sorted from pLN of mice immunized 5.5 days earlier with ovalbumin (OVA)/Montanide (in case of macrophages GM-CSF was added) (n≥3). Data shown in A and B represent two independent experiments (n=4). Data shown in D are representative of three independent experiments. Bar graphs and scatter dot plots show the mean ± STD. Statistics in (B) using unpaired t-test. \**P <* 0.05, \*\**P <* 0.005 and \*\*\**P <* 0.001

### FRC express the prostanoid PGE_2_ which acts on EP2/4 receptors on T cells to inhibit their expansion

To investigate under which conditions the iNOS and COX pathways become active in lymph node fibroblasts, we initially analyzed the transcript levels of these enzymes in pLN2 cells as well as ex vivo LN FRC. Consistent with previous data (9-11) the gene coding for iNOS, *nos2*, was not detectable in naive pLN2 but strongly induced 7h after stimulation with IFNγ/TNFα or LPS while *ptgs2* (COX2) was highly expressed in unstimulated pLN2 without major changes upon cytokine stimulation (Supplementary Fig. 1B). Similarly, *ptgs1* (COX1) transcripts were constitutively expressed in pLN2 but at 10-fold lower levels than *ptgs2* suggesting COX2 is the isoform preferentially expressed in pLN2 cells and responsible for the attenuating effect observed on T cell expansion. To test whether this holds true *in vivo*, peripheral LN (pLN) and spleen from adult wt mice were passed through a mesh to enrich either for lymphocytes or ‘non-soluble’ stromal cells (4). Transcripts for both COX isoforms were enriched 5-50 fold in the stromal cell fraction of pLN with less marked differences observed for the splenic fractions (Supplementary Fig.1C). To identify the stromal cell types expressing *ptgs1/2* in naïve pLN, various cell populations from naïve pLN were purified and analyzed. While *ptgs1* transcripts were found at similar levels in FRC and endothelial cells, *ptgs2* transcripts were expressed at 100-fold higher levels in FRC relative to endothelial, myeloid or lymphoid cells (Fig.1C). Notably, transcript levels of *ptgs2* were approximately 1000-fold higher in FRC than of *ptgs1* indicating COX2 is the principle isoform expressed by naïve pLN FRC.

COX enzymes are critical enzymes for the production of lipid intermediates that get further metabolized to produce the different effector molecules of the prostaglandin and thromboxane family. Previous studies have shown that high COX2 expression often correlates with high levels of PGE_2_, with this prostanoid being known to inhibit T cell activation and proliferation in various settings (12-16). As PGE_2_ production depends on prostaglandin E synthases, we assessed the transcript levels of all three isoforms (*ptges1-3*). Both in naïve pLN and spleen all three isoforms were expressed in the stromal cell fraction with *ptges1 and ptges3* being preferentially expressed in that fraction (Supplementary Fig. 1D). Consistent with a constitutive activity of this pathway in FRC, overnight culture of the pLN2 FRC cell line led to the accumulation of 10-15ng/ml PGE_2_ in the supernatant which was abolished in presence of indomethacin (Fig. 1D). In addition, pLN FRC isolated and sorted from d5 OVA/Montanide immunized mice produced high concentrations of PGE_2_ as well, about 10-fold more than macrophages isolated from the same LN (Fig. 1E). These findings point to FRC being a rich constitutive PGE_2_ source in naïve and activated pLN.

To assess the direct effect of PGE_2_ on T cell activation and proliferation, naïve T cells were stimulated in the presence of increasing PGE_2_ concentrations. In line with previous data (12) we found that the proliferation of CD8+ and to a greater extent CD4+ T cells was impaired in presence of PGE_2_ (Fig. 2A) at concentrations similar to those produced by FRC. Dampened T cell proliferation was accompanied by reduced CD25 upregulation and reduced CD62L loss on total and proliferating T cells whereas CD44 levels were less altered (Fig. 2B and Supplementary Fig. 2A, B). To determine if PGE_2_ is the attenuating factor produced by COX2-expressing FRC we used antagonists for the two high affinity receptors for PGE_2_, EP2 and EP4, expressed by T cells (12, 14). Indeed, CD4+ and CD8+ T cell proliferation was rescued when the EP4 inhibitor was added to the co-culture system, to an extent comparable to indomethacin itself (Fig. 2C, D). The effect of the EP2 inhibitor was less pronounced. In summary, these *in vitro* results support the notion that PGE_2_ may be the major COX2 dependent prostanoid released by pLN FRC dampening T cell responses by signaling via the EP4-receptor on naïve T cells.

**Figure 2:**
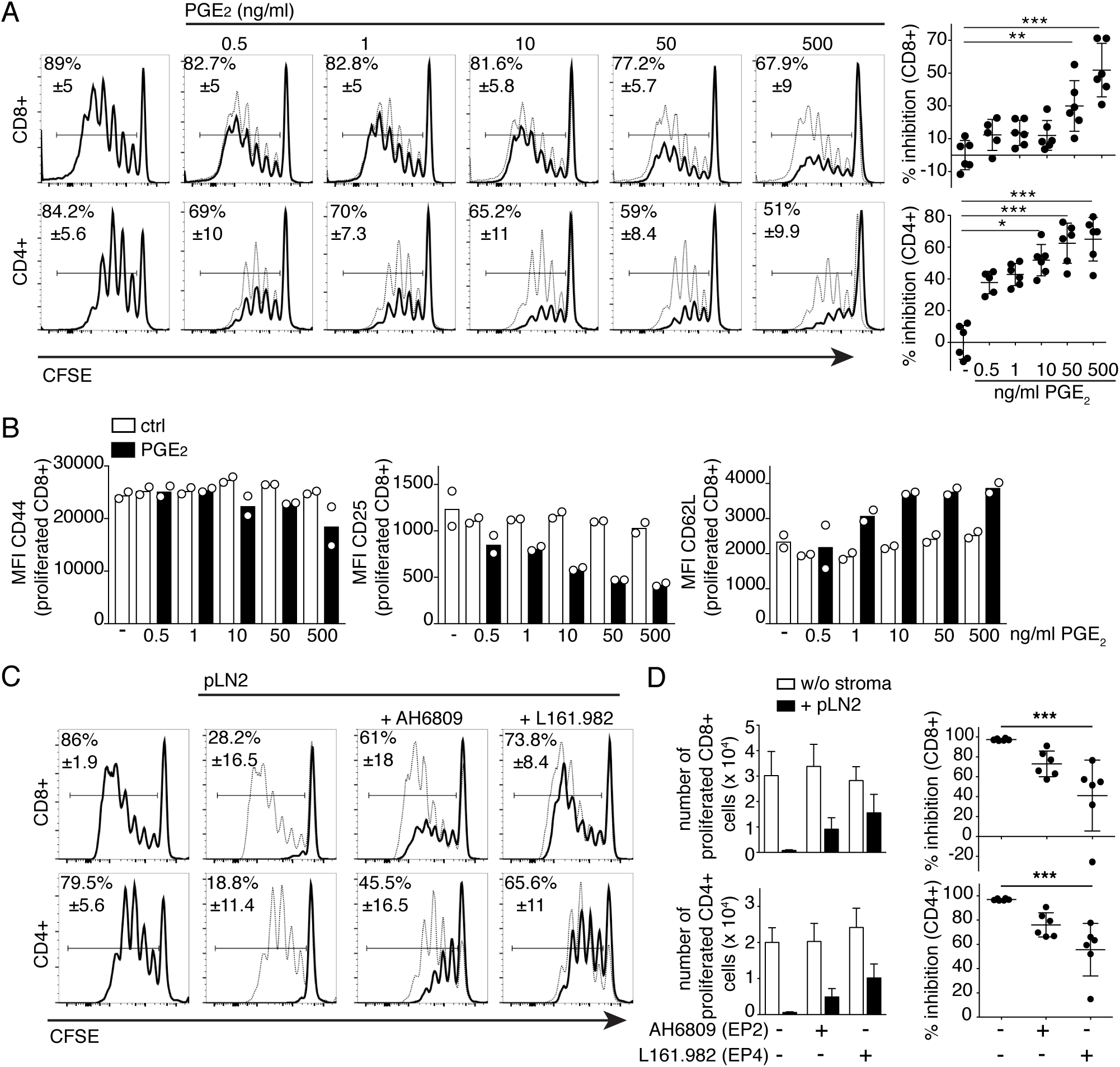
LN FRC constitutively produce the COX-dependent mediator PGE_2_ that enables inhibition of T cell activation and proliferation via EP4 receptor signaling. Flow cytometric analysis of CFSE dilution in CD8+ and CD4+ T cells that were activated with αCD3/28 DynaBeads and cultured for 3d. (A) Left side: CFSE dilution profiles of T cells in presence of the indicated concentrations of PGE_2_ (black line) or solvent control (Ethanol; thin line), with on the right side scatter plots showing the percentage inhibition of T cell proliferation. (B) Expression level (MFI) of CD44, CD25 and CD62L on proliferated CD8+ T cells in presence of PGE_2_. (C) CFSE profile of T cells cultured in presence (thick line) or absence (thin line) of pLN2 cells and the EP2 antagonist AH6809 (5µM), or the EP4 antagonist L161.982 (5µM). (D) Number of proliferated T cells (left side) and percentage inhibition of T cell expansion (right side) as in (C). (A-D) Data are representative of 3 independent experiments (n=6). All bar graphs and scatter plots show mean ± STD. \**P <* 0.05, \*\**P <* 0.005 and \*\*\**P <* 0.001 (Kruskal Wallis followed by Dunns post-test).

### FRC-derived prostanoids can suppress both weak and strong T cell responses *in vitro* while FRC-derived NO mainly dampens strong T cell responses

Given that we have identified COX2/PGE_2_ as the second pathway in FRC able to attenuate T cell responses, besides iNOS/NO, we wished to address why FRC have two distinct mechanisms to dampen T cell expansion. Based on the constitutive versus inflammation-induced expression of COX2 and iNOS, respectively, we hypothesized that iNOS-mediated T cell inhibition may only act during strong T cell responses, when a certain threshold level of inflammation is reached, whereas COX2 mediated T cell inhibition may act on strong and weak T cell responses and thus affect also low affinity and possibly autoreactive T cells. To test this hypothesis, we used various model systems to mimic strong versus weak T cell responses. First, we analyzed OT-1 T cell proliferation upon stimulation with BM-DCs loaded with different concentrations of ovalbumin (OVA) peptide, either of high (N4) or low (V4) affinity for the OT-1 T cell receptor (TCR) (21). Although at least 100 times more V4 than N4 peptide was needed to induce T cell proliferation and CD44 upregulation *in vitro*, we did not find a suitable condition leading to weak T cell expansion (Supplementary Fig. 3A), in line with previous evidence (21). When FRC were added in a setting of strong TCR stimulation, due to a high N4 concentration, or in settings of weaker TCR stimulation, such as low N4 or high V4 concentration, always a robust inhibition of T cell proliferation was observed (Supplementary Fig. 3B). Relatively high NO levels were found in the supernatant of all three conditions indicating that there was sufficient inflammation to induce iNOS expression (Supplementary Fig. 3C). As BMDC may contribute NO or Cox-dependent prostanoids in this assay (Fig.S3 C) (11, 16), we decided to investigate different strengths of T cell activation in the absence of DC, by using beads loaded with different concentrations of αCD3/28 antibodies. In this experimental system, increasing αCD3/28 concentrations were reflected by a more gradual increase in the expansion of CD4+ and CD8+ T cells (Fig.3 A and B). Adding pLN2 cells in lower or higher numbers attenuated the T cell expansion more extensively with weaker than stronger T cell stimuli, especially for CD8+ T cells, with this effect being also visible at the level of CD25 and CD44 upregulation (Fig.3 A and B). NO levels increased with increasing FRC number and TCR strength, but showing an inverse correlation with T cell inhibition by FRC (Fig. 3A, C). In previous work, early IFNγ by T cells was shown to be responsible for iNOS/NO expression in FRC, which then acts in a negative feedback loop to reduce the number of T cells recruited into the response (9-11). When cocultures of FRC and T cells were analyzed already on d1 of co-culture a clear correlation was observed between the strength of TCR activation and the percentage of activated and IFNγ + CD8+ T cells induced, as well as with the NO level in the culture supernatant (Supplementary Fig.4 A-C). Flow cytometric and histological analysis confirmed the correlation between iNOS protein expression in FRC with TCR signal strength and IFNγ release (Fig.4 A and B). Indeed, pharmacological inhibition of iNOS activity with 1400W abolished the suppressive effect by FRCs in the setting of a strong but not weak T cell stimulation (Fig. 4C). These findings support our hypothesis that only strong T cell stimulation leads to IFNγ secretion by T cells sufficient to induce NO in FRC which then limits the number of T cells recruited into the response.

**Figure 3:**
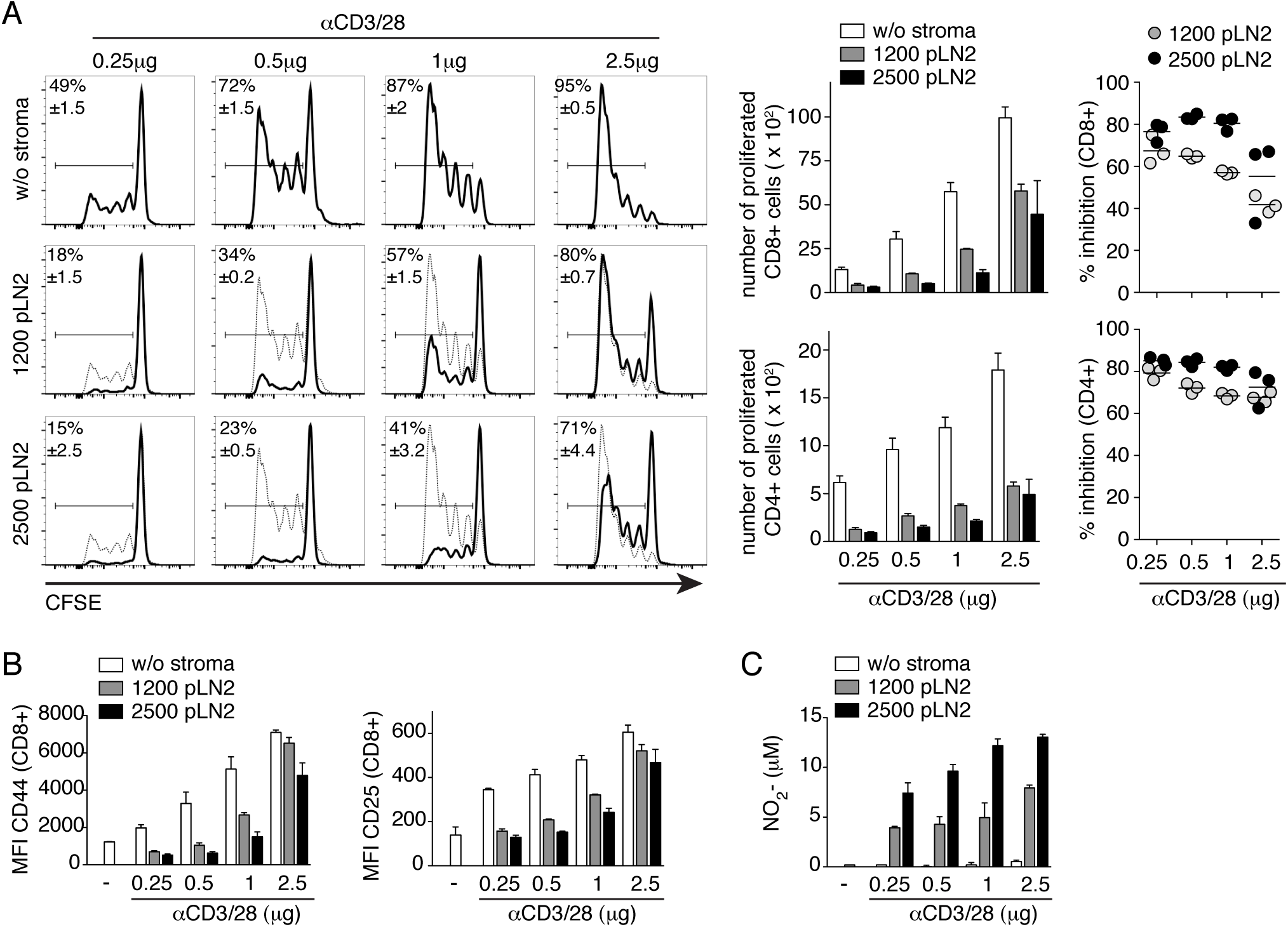
FRC can dampen T cell responses of various strengths. Flow cytometric analysis of CFSE dilution in CD8+ and CD4+ T cells that were activated with beads coated with indicated concentrations of αCD3/28 and cultured for 3d ± the indicated numbers of pLN2. (A) Left side: FACS analysis of CFSE-labeled CD8+ T cells in presence (black line) or absence (thin line) of FRC. In the middle: Bar graphs showing the number of proliferated CD8+ and CD4+ T cells. Right side: Scatter plots showing the percentage inhibition of T cell expansion by FRC. (B) Expression level (MFI) of CD44 and CD25 on CD8+ T cells. (C) Griess assay measuring nitrite levels as surrogate for NO levels in the supernatant of T cell-FRC co-cultures described in A. (A-C) Data are representative of 4-5 independent experiments with three replicates each. All bar graphs and scatter plots show mean ± STD.

Previous evidence for a role of iNOS in dampening T cell expansion *in vivo* have been collected for strong immune responses only (9-11). To test whether this process is absent for weaker immune responses, wt and iNOS^−/−^ mice having received ovalbumin-reactive CD8+ (OT-1) T cells were immunized s.c. with different doses of OVA/Montanide to mimic different TCR stimulation levels (Fig. 4D). While on average there was an increased OT-1 T cell response in iNOS*-*deficient mice that received high antigen doses compared to wt control mice, this difference was not observed with low or intermediate antigen doses. Similar to previous studies (10, 11), no differences were observed in the expression of activation marker or in the killing capacity of OT-1 T cells isolated from either wt or iNOS*-*deficient mice (Supplementary Fig.4 D and E).

**Figure 4:**
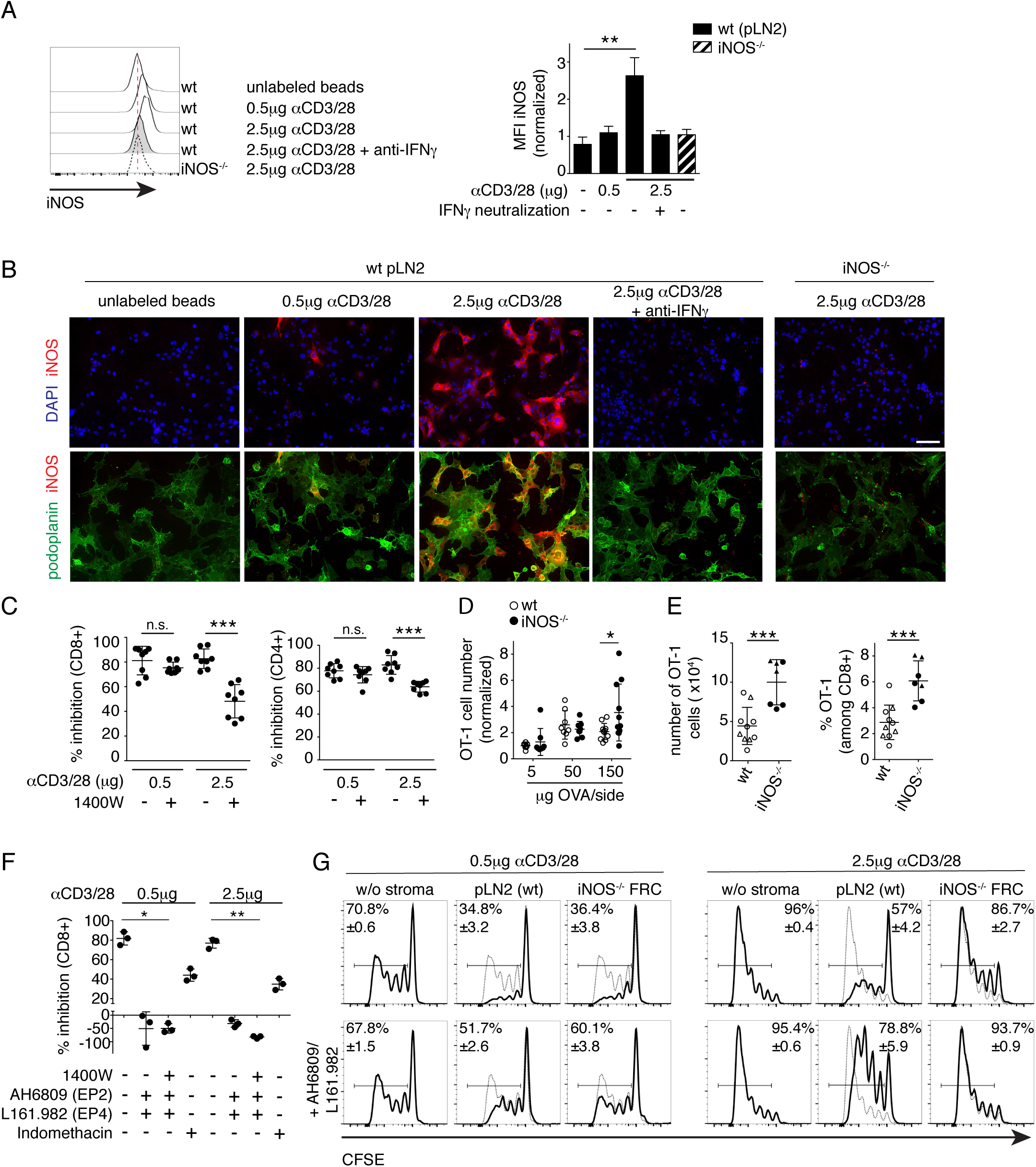
iNOS mediated T cell inhibition by FRC arises during strong but not weak T cell responses. Flow cytometric (A, C) and histological analysis (B) of 2 days cocultures containing FRC and CD8+ and CD4+ T cells activated with beads coated with the indicated dose of αCD3/28. (A) Intracellular iNOS protein expression in pLN FRC cell lines of wt (pLN2) or iNOS*^−/−^* origin, co-cultured for 2d with T cells ± anti-IFNγ (10µg/ml). Histograms showing flow cytometric detection (A; n=5, pool of 3 independent experiments) and photos depicting histological detection of iNOS (red), along with a nuclear marker (DAPI, blue) or a FRC marker (podoplanin, green) (B). Scale bar, 100µm; representative images of 3 independent experiments. (C) Activated T cells cultured for 3d ± pLN2 ± iNOS inhibitor 1400W (3µM) and assessed for the percentage inhibition of T cell expansion (n=8, pool of 3 independent experiments). (D-E) Wt and iNOS*^−/−^* mice received OT-1 CD8+ T cells i.v. followed by. s.c. immunization with the indicated concentrations of OVA/Montanide/Poly(I:C), with draining LNs investigated by flow cytometry. (D) Scatter plot shows OT-1 T cell number per dLN of wt versus iNOS*^−/−^* mice on d4 after immunization. Pool of 2-3 independent experiments (n≥8), with data normalized to the average number obtained with the 5µg OVA condition. (E) On d37 after primary immunization with 50ug OVA/Montanide with either 25µg (triangles) or 10µg (circles) Poly(I:C) s.c., mice were re-challenged with 5µg OVA/Montanide/Poly(I:C) s.c. and dLN investigated 3d later. Bar graphs depict numbers and percentages of OT-1 cells in wt or iNOS*^−/−^* mice, respectively (n≥7; pool of two independent experiments). (F-G) Flow cytometric analysis of T cell expansion activated with αCD3/28 coated beads and cultured for 3d in the presence or absence of FRC. (F) Scatter plot depicting the percentage inhibition of CD8+ T cell proliferation in presence of the indicated inhibitors: 1400W (for iNOS; 3µM), AH6809 (for EP2; 5µM); L161.982 (for EP4; 5µM); indomethacin (for Cox2; 10µM) (n=3; data representative of 3 independent experiments). (G) CFSE profiles of CD8+ T cells in the presence (thick line) or absence (thin line) of wt pLN2 or iNOS *-/-* FRC ± AH6809/L161.982 (5µM each). Data are representative of 3 independent experiments with 2-3 replicates each. Bar graphs and scatter plots showing mean ± STD. Statistics: (A and F) Kruskal Wallis followed by Dunns post-test (C-E) unpaired t-test; \**P <* 0.05, \*\**P <* 0.005 and \*\*\**P <* 0.001

Having shown that iNOS expression in FRC depends on the level of IFNγ secreted by the neighboring T cells, we hypothesized that the suppressive effect mediated by iNOS may be more pronounced in a secondary response by memory T cells able to sense less antigen and produce IFNγ more quickly (22, 23). To this end, wt and iNOS*^−/−^* mice immunized with low or intermediate doses of OVA/Mont were rechallenged after 5 weeks with a low dose of OVA/Mont to measure the OT-1 memory cell response. Indeed, the frequency and numbers of OT-1 T cells were significantly higher in draining pLN of iNOS*^−/−^* compared to wt mice (Fig. 4E). Together, these findings suggest that FRC sense the strength of the T cell response and produce NO only in the case of a strong primary T cell response. Interestingly, this inhibitory feedback loop appears to be more pronounced for secondary T cell responses, as it extends then also to lower antigen doses, thereby limiting the size of the effector T cell pool generated.

With iNOS being induced in FRC during strong primary T cell responses, we wished to investigate if the constitutive COX2 pathway may have a complementary role by inhibiting both the strong and weak T cell responses. Co-culture studies revealed an inhibitory role for COX and PGE_2_-receptors EP2/4 in weak but also strong CD8+ and CD4+ T cell responses (Fig.4F and Supplementary Fig.4F). This was confirmed by using a newly generated iNOS*^−/−^* FRC cell line that did efficiently impair the expansion of weakly but not strongly stimulated T cells (Fig. 4G). Also in this experimental setup, addition of EP2/4 antagonists resulted in the rescue of both, weak and strong T cell responses, suggesting that the COX2/PGE_2_ pathway can dampen T cell responses independently of the signal strength.

### FRC-specific deletion of COX2 does not affect T cell homeostasis or clonal expansion in vivo

As *nos2^flox/flox^* mice have not yet been reported, the specific *in vivo* role of iNOS in LN FRC relative to other cell types cannot be studied yet. In contrast, *ptgs2^flox/flox^* mice have been generated (24). Therefore, we made mice deficient in COX2 within FRC by intercrossing *ccl19cre* with *ptgs2^flox/flox^* mice (COX2^ΔCCL19cre^) to test whether LN FRC can indeed dampen T cell responses *in vivo* via COX2. First, we measured the specificity and efficiency of Cre recombinase-mediated EYFP expression within different LN cell populations using COX2^ΔCCL19cre^ mice crossed with the Cre-reporter strain *ROSA26^eyfp^* (ROSA26-EYFP^CCL19Cre^) Flow cytometric analysis revealed that 90% of pLN FRC were EYFP+, whereas less than 15% of endothelial cells and CD31-pdpn- cells were EYFP+, with no detectable Cre activity observed within hematopoietic cells (Supplementary Fig. 5A and data not shown). Among the two major FRC subsets, an average of 96% of T zone FRC and 76% of Medullary FRC exhibited Cre activity in naïve LN. This is in line with COX2^ΔCCL19Cre^ mice completely lacking *ptgs2* mRNA expression in T zone FRC (Fig. 5A). Ablation of Cox2 expression did neither alter *ptgs1* transcript levels nor affect *ccl19* expression in these cells (data not shown). Given that COX2/PGE_2_ expression is constitutive in LN FRC, we analyzed the hematopoietic and non-hematopoietic cell types in spleen and pLN of naïve COX2^ΔCCL19cre^ mice using flow cytometry and histology, however, no major difference was observed in the size and organization of lymphocyte and FRC populations (Fig. 5B-D and Supplementary Fig.5B and C). Further, the activation status of T cells found in naïve mice was comparable between COX2^ΔCCL19cre^ and control mice, suggesting that Cox2 in FRC is not required to regulate T cell homeostasis. To assess whether COX2 activity in FRC plays a role in regulating T cell expansion in response to foreign antigen, we infected COX2^ΔCCL19cre^ and control mice with a high dose of lymphocytic choriomeningitis virus (LCMV) clone 13 that establishes a chronic non-resolving infection. This model system allows to look at the initial T cell expansion phase in addition to studying the chronic phase of a viral infection. Notably, PGE_2_ has been recently proposed to inhibit chronic T cell responses to clone 13 infection (18). Infected COX2^ΔCCL19cre^ mice showed an expansion of LCMV-specific CD8+ T cells within pLN and spleen that was comparable to wt mice on day 8 p.i. (Fig. 5E; Supplementary Fig. 5D and E). Similar results were obtained for the peak of the T cell response in pLN draining the site of s.c. vaccination with OVA/Montanide on d3 and d5 (data not shown) suggesting a redundant function for COX2 in FRC during the T cell expansion phase. Notably, LCMV-specific CD8+ T cells found in LN and spleen of d8 infected COX2^ΔCCL19cre^ mice showed a higher surface expression of PD-1 (Fig. 5F and Supplementary Fig. 5F), which is a negative regulator of T cell activation in chronic T cell responses to persisting infections or tumors (25, 26). Similar findings have been reported recently for LCMV-specific CD8+ T cell responses in mPges1- or EP2/4-deficient mice (18) suggesting that this effect is most probably PGE_2_ driven. To assess effector function of LCMV specific CD8+ T cells in these two mouse models on d8 p.i., we determined their cytokine production as well as the viral clearance by measuring the virus titer in the blood. Indeed, CD8+ T cells isolated from COX2^ΔCCL19cre^ mice exhibited a slightly reduced IFNγ and TNFα production upon re-stimulation with different LCMV peptides (Supplementary Fig. 5G), suggesting COX2 in FRC has weak proinflammatory effects on the early phase of the T cell response. However, the viral burden in the blood was similar in both groups (Fig. 5G). In line with previous studies (27, 28), our histological analysis showed a dramatic impact of LCMV infection on the LN compartmentalization as well as the function of LN stroma cells as CCL21 source (Supplementary Fig. 5H) which was comparable between COX2^ΔCCL19cre^ and control mice. This defect was transient in both mouse models as the lymph node organization and CCL21 expression was restored by day 21 post infection (Fig. 6A).

**Figure 5:**
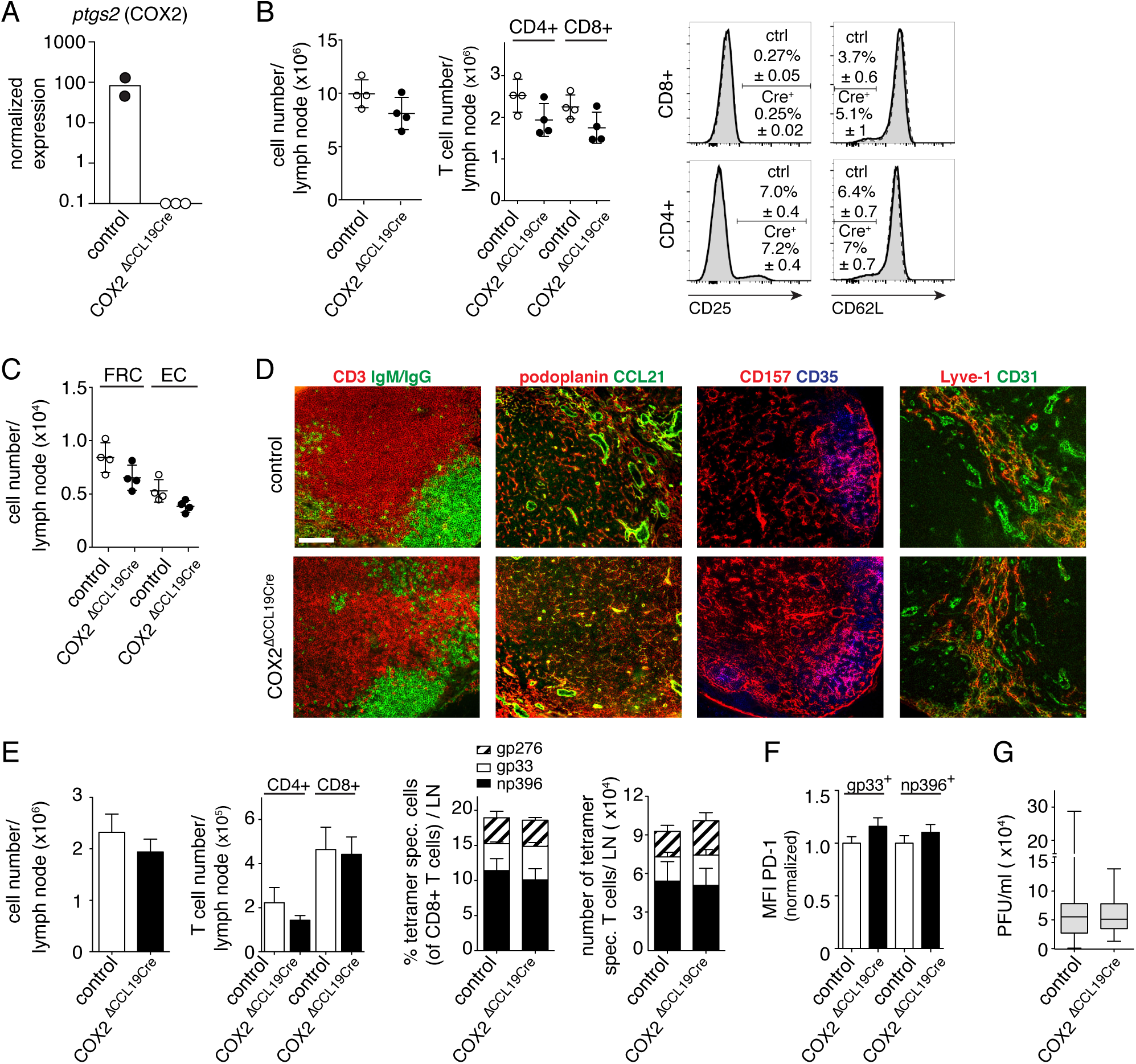
Deletion of COX2 specifically in FRC does not alter T cell homeostasis or peripheral LN structure. Characterization of pLN of mice genetically lacking COX2 expression specifically in FRC (COX2^ΔCCL19Cre^) versus their littermate wt mice (called ‘controls’). (A) Transcript levels of *ptgs2* were analyzed in sorted T zone FRC (TRC; CD45- CD35- CD31- Podoplanin+ CD157+) isolated from naive pLN of Cre+ vs Cre- mice (n=2-3, each sample represents a pool of 2-3 mice). Transcript levels below the detection limit or non-specific transcripts are indicated as white circles on the x-axis. (B) Scatter plots showing total cell numbers (left side) or CD4+ and CD8+ T cell numbers (middle) in digested naive pLN of the indicated mice, or representative histograms (right side) showing the percentage of activated CD8+ and CD4+ T cells, by gating on CD25^high^ or CD62L^low^ cells, in samples derived from Cre+ (thick line) or Cre- littermates (dashed line with gray shading). Bar graphs and scatter plots showing mean ± STD. (C) Scatter plots showing total FRC and endothelial cell (EC) numbers per naïve pLN of the indicated mice (B-C: n=4; representative of three independent experiments). (D) Immunofluorescence microscopy analysis of labeled pLN sections of naïve Cre+ mice versus controls. Localization of T and B cells as well as antibody staining for different stromal cell types or their products are shown. Data are representative for 2 independent experiments investigating 3 mice per genotype. Scale bar, 100µm. (E-G) Cre+ and Cre- mice were infected i.v. with LCMV clone 13 with analysis performed at 8d post infection, using flow cytometry (E, F) or plaque assay (G) (pool of two independent experiments; n≥7). (E) Bar graphs showing total cell numbers and T cell numbers per digested pLN, as well as LCMV-specific CD8+ T cells in percentage and numbers (for the three indicated tetramer specificities). (F) MFI of PD-1 expression on gp33- versus np396-specific CD8+ T cells, with results from Cre+ mice normalized to control mice. (E-F) Bar graphs showing mean ± SEM. (G) Box plot showing viral titers in the blood of the indicated mice.

### FRC-specific deletion of COX2 leads to a stronger T cell response in the chronic phase of LCMV infection and a better virus control

Given the higher T cell responses previously observed in mPges1^−/−^ mice (18), we wished to assess whether COX2 expression in FRC was responsible for suppressing chronic T cell responses. First, we confirmed that LN FRC are the principal cell type displaying Cre-activity on d19 p.i. with clone 13 virus (Supplementary Fig. 6A and data not shown), similar to naive mice. On d21 post infection, we detected a significant increase in number of total and LCMV-specific CD8+ T cells in pLN and to a lesser extent in the spleen of COX2^ΔCCL19cre^ relative to wt mice, especially for np396-specificities (Fig. 6B, Supplementary Fig. 6B). While the LCMV-specific cells isolated from COX2^ΔCCL19cre^ mice showed a similar PD-1 and IFNγ expression level as control mice, the TNFα expression level was increased in cells isolated from pLN (Fig. 6C and D) but not the spleen (Supplementary Fig. 6C). Given the strong increase in gp33- and np396-specific T cells in LN of d21, the total number of IFNγ+ and TNFα+ T cells is increased by more than 100% in COX2^ΔCCL19cre^ mice (Fig. 6E). This increased CTL response translated into a significant improvement in viral clearance in the blood and LN of COX2 deficient animals (Fig. 6F and data not shown). Together these findings demonstrate that COX2 activity within LN FRC has an important role in suppressing chronic T cell responses during persisting viral infections, by downregulating CTL numbers and function and consequent reduction of viral clearance.

**Figure 6:**
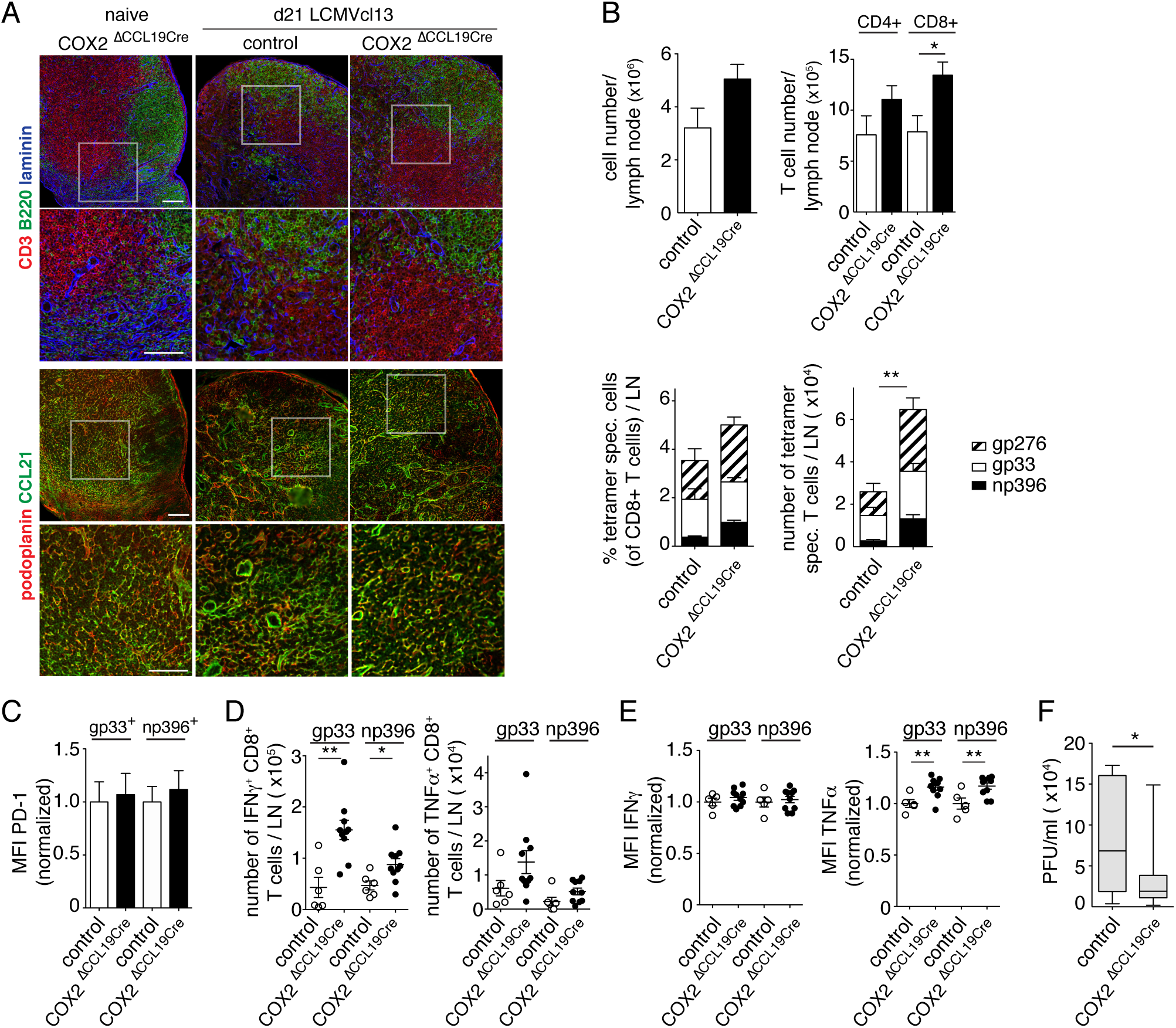
COX2 activity in FRC dampens chronic T cell responses during persisting LCMV infection. COX2^ΔCCL19Cre^and control mice were infected with LCMV clone 13 and analysis performed at 21d post infection, using histology (A), flow cytometry (B-E), and plaque assay (F). (A) Histological analysis of pLN for the indicated lymphocyte and fibroblast markers. Squares indicate the regions shown enlarged in the row below. Images are representative for 3 mice/genotype. Scale bar, 100µm. (B) Bar graphs showing total cell numbers and T cell numbers per pLN, as well as the percentage and number of LCMV-specific CD8+ T cells in the indicated mice. (C) PD-1 expression levels (MFI) on gp33- versus np396-specific CD8+ T cells from Cre+ mice, normalized to control mice (B and C: data show a pool of at least 3 independent experiments; n ≥6). Numbers (D) and MFI (E) of IFNγ and TNFα expression in LCMV-specific CD8+ T cells in pLN after re-stimulation with gp33 or np396 peptides (pool of two independent experiments; n≥5). (F) Viral titers in the blood of the indicated mice (pool of blood samples from 5 independent experiments; n≥ 10). (B-E) Bar graphs showing mean ±SEM. Statistics: Unpaired t-test; (normally distributed data) or Mann-Whitney test (for not normally distributed data). \**P <* 0.05, \*\**P <* 0.005 and \*\*\**P <* 0.001.

## Discussion

In the current study, we show that FRC display a highly active COX2/PGE_2_ pathway that dampens the expansion of *in vitro* activated CD8 and CD4 T cells by acting via EP2/4 receptors. This pathway dampens both weak and strong T cell responses, in contrast to FRC-derived NO which is produced mainly during strong TCR stimulations pointing to two distinct inhibitory pathways in FRC. Using mice selectively lacking COX2 in lymphoid tissue FRC we provide direct evidence for an *in vivo* function of FRC in downregulating T cell responses, not during the early phase of an acute CD8 T cell response but later during a chronic response. Thus, pLN FRC and COX2 expressed by them can have anti-inflammatory roles on T cell responses in vivo.

FRC have been proposed to play a dual role for adaptive immunity, either promoting or inhibiting it (1, 2), similar to myeloid and T cells. *In vitro* experiments have revealed that FRC can act as bystander cells to dampen T cell responses, independently of effects on regulatory T cells or APC, by sensing IFNγ and TNFα released early during T cell priming and leading to FRC-derived NO release (9-11). This latter concept is consistent with *in vivo* evidence of an exaggerated T cell response to immunogenic stimuli in iNOS-deficient mice (9-11). We now extend these findings by showing that iNOS expression and NO release by FRC *in vitro* is mainly observed during strong T cell responses characterized by a marked early IFNγ secretion, but not in weaker T cell responses where IFNγ levels are low. Due to the lack of iNOS^fl/fl^ or suitable chimeric mice (9, 11) this concept could only be tested using iNOS^−/−^ mice where myeloid cells are also iNOS-deficient. Consistent with our *in vitro* findings, we show here that primary T cell responses were enhanced in iNOS^−/−^ mice only in response to a high but not low antigenic stimulation. More striking is the enhancement we observed for the secondary T cell expansion to a low antigen dose in iNOS^−/−^ mice, similar to an earlier report (23) and consistent with an expected enhancement of the early IFNγ expression by restimulated memory T cells (22). Therefore, this regulatory pathway via NO release may only come into effect in acute type I immune responses which have the potential to damage neighboring cells. An interesting application of this property of LN FRC or other mesenchymal stromal/stem cells (MSC) is their use in settings of acute inflammation, such as transplantation or sepsis, where proinflammatory cytokines are abundant and fibroblasts can unleash their full inhibitory and tissue-preserving function, at least in part by releasing NO (29, 30). However, both MSC and LN FRC have alternative ways of inhibiting T cell responses as they can express PD-L1, TGFβ, COX2/PGE_2_ and possibly other factors (11, 29, 31-34). Here we provide detailed evidence for a second inhibitory pathway in LN FRC, which is still active in iNOS^−/−^ FRC and is mediated by the COX2-dependent synthesis of PGE_2_.

We tested the importance of the COX2 pathway specifically in FRC, both *in vitro* and *in vivo*. We observed that COX2 is expressed constitutively in FRC of naïve LN at levels clearly above those observed for COX1, consistent with a recent report (31). This is opposite of the usual COX expression pattern, with COX1 being typically constitutively expressed and COX2 being induced during inflammation (12, 16). LN FRC maintain this property even *in vitro*, including after several weeks of culture, suggesting this is an intrinsic and imprinted feature in murine FRC (11, 31). This expression is not much altered upon stimulation by cytokines or TLR2/4 ligands (11), in contrast to most other cell types, including myeloid cells, that depend upon these signals for COX2 expression (16). Of note, the gut lamina propria is one of the rare tissues displaying high COX2 expression and thereby an immune microenvironment that attenuates T cell immunity to the diverse commensals and food antigens. Interestingly, adherent stromal cells were identified as major source of COX2, displaying constitutive PGE_2_ expression without need for exogenous stimuli (35, 36). A recent study revealed that also human FRC are capable to suppress T cell proliferation and differentiation in a COX-dependent manner (34). In contrast to murine FRC, both COX1 and COX2 appear to be constitutively expressed in human FRC (31, 34).

COX enzymes are the rate-limiting enzymes for the generation of five different prostanoids, including PGE_2_. We decided to focus on this lipid mediator, as it is a known negative regulator of T cell responses, and as its high constitutive expression in LN FRC came as a surprise to us given that LN are sites of adaptive immune response induction. Remarkably, the expression in FRC was 50-fold above LN macrophages for COX2 transcripts and 10-fold for PGE_2_ suggesting FRC may be poised to dampen T cell responses. Indeed, ex vivo FRC inhibited CD4 and CD8 T cell activation and expansion in a COX2-dependent manner, and to an extent similar to purified PGE_2_ (31). In addition, blocking of the two major PGE_2_-receptors on T cells, EP2 and EP4, mimicked this effect, providing strong evidence for PGE_2_ being the principal COX2-dependent prostanoid responsible for attenuating T cell responses in co-cultures with pLN FRC.

Given the high expression of COX2 and PGE_2_ in pLN FRC before the start of T cell priming, we hypothesized this pathway could ensure that responses by non-specific or weakly activated T cells are avoided (2). When we varied either the antigen affinity or quantity in the co-culture assay, FRC-mediated suppression abolished responses to weaker stimuli, while stronger stimuli overcame these inhibitory effects despite the combined presence of COX2- and iNOS-dependent factors. Importantly, inhibition of COX2 or EP2/4 but not iNOS allowed again the expansion of weakly responding T cells in FRC containing cultures. Several findings support the notion that less T cells got recruited into the response: we observed 1) more undivided cells; 2) a reduced CFSE dilution of those cells that did divide; 3) diminished expression of activation markers on undivided but not proliferating T cells; 4) strongly reduced numbers of antigen-specific T cells generated; 5) fewer effector cells capable of expressing IFNγ while killing of targets occurred normally. These observations are consistent with earlier reports on PGE_2_ effects on T cells, both by interfering with TCR signaling as well as by deviating in some settings from an IFNγ-driven response (12-14). For example, when human CD4+ T cells were activated by suboptimal anti-CD3/CD28 stimulation *in vitro*, most transcriptional changes were ablated by a simultaneous PGE_2_ exposure, leading to strongly reduced cell cycle entry while not altering proapoptotic pathways (20). Lck, zap70 and Ca-flux have been identified as the main targets of negative regulation by PGE_2_ in T cells, with the suppression being surmountable by a stronger stimulation (20). These findings were recently confirmed in the context of murine T cells cocultured with either murine or human LN FRC (31). Interestingly, NO appears to mediate its inhibitory effect also by interfering with early TCR signaling, such as by nitrosylation of CD3-zeta (37) thereby blocking its phosphorylation sites needed for cell activation. While there is evidence for reciprocal regulation between the iNOS and COX2 pathways (38, 39), the observation that iNOS^−/−^ as well as COX2^−/−^ FRC (31) can still suppress via the other pathway suggest that at least part of their T cell inhibitory function is independent. This notion is further supported by the COX2-dependent inhibition of weaker T cell responses where iNOS expression is very low. In conclusion, while NO expression by FRC is transient and limited to strong type I T cell responses, COX2/PGE_2_ expression in LN FRC is constitutive and can affect T cells throughout the response and independently of signal strength.

Our findings based on co-culture assays raise the question of *in vivo* relevance, namely do COX2-dependent mediators derived from FRC inhibit T cell responses in settings of homeostasis, vaccination or disease. On one hand, FRC may participate in maintaining peripheral tolerance by ablating the typically low affinity responses to self-antigens. So far, we have not observed any obvious autoimmune phenotype in mice lacking COX2 in FRC (COX2^ΔCCL19cre^) or in all body cells with mice aged for 20 weeks in our SPF facility (data not shown). This is consistent with previous observations in COX2^−/−^ mice (16, 40) as well as with people treated over months or years with COX-inhibitors where autoimmune side-effects have rarely been reported. On the other hand, we hypothesized that FRC-derived PGE_2_ may set a threshold for T cell activation in response to foreign antigens either to prevent unnecessary T cell activation in case of very weak inflammatory stimuli, or to focus the response to higher affinity T cells. Our approach to vaccinate with OVA/Montanide in wt versus COX2^ΔCCL19cre^ mice did not show a marked change in the early expansion of high affinity CD4+ OT-2 and CD8+ OT-1 T cells (data not shown), or of LCMV-specific CD8+ T cells at the peak of infection. This is in contrast to Cui and colleagues that recently reported an increased OT-1 T cell expansion at 48h upon vaccination with OVA-loaded BMDC in mice lacking an enzymatically active COX2 (COX2^Y385F/Y385F^); however, this difference was lost 24h later (31). Currently, the reason for this discrepancy is unclear but could be based on differences in timing or antigen dose reaching the LN. It seems also plausible that COX2- dependent processes in non-FRC may have contributed to the phenotype described by Cui and colleagues as activated macrophages or DC were also deficient in prostanoid synthesis in their mouse model (31). Of note, they observed only a delay of the response and not a qualitative or quantitative difference at time points later than 48h. Nevertheless, these data indicate that PGE_2_ present within the LN may modulate in some settings the T cell priming or early expansion phase, and possibly clonal selection and amplification.

Given the normal T cell expansion we observed in the early phase of the response to LCMV we focused our attention on the chronic phase where mice deficient in EP2/4-expression in activated T cells or globally deficient in mPges-1 have previously been shown to have an enhanced antiviral CD8+ T cell response on d21 post LCMV infection, both in number, effector function and reduction of the ‘exhaustion’ phenotype (18). That study demonstrated that PGE_2_ acts on virus-specific CD8+ T cells via EP2/4 and restricts their survival but not their proliferation, presumably without Treg involvement. Interestingly, EP2/4 expression is increased in PD-1^hi^ T cells on d21, and inhibiting both COX2 and PD-L1 showed additive effects on T cell responsiveness (18). However, this report did not reveal which cells were the main targets of COX2 inhibition. Interestingly, we observed that COX2^ΔCCL19cre^ mice reproduced the findings of Kaech and colleagues, notably the increased numbers of endogenous virus-specific CD8+ T cells of three different specificities, along with their effector function leading to a better virus control without evidence for increased immunopathology (18). Therefore, we propose that FRC within T zones of SLO, like LN and spleen, negatively regulate the survival of chronically activated CD8+ T cells via their constitutive production of PGE_2_, presumably by enhancing the chronic TCR signals that drive T cell exhaustion. Importantly, COX2 inhibition in FRC can partially reverse this effect.

COX inhibitors are the most frequently used drugs to prevent inflammation. Therefore, our results suggesting COX2/PGE_2_-expressing FRC lead to an inhibition rather than enhancement of T cell responses may seem paradoxical. However, COX-dependent prostanoids and PGE_2_ in particular are well known for their dual role in inflammation and immunity (12-15). PGE_2_ has been observed to be anti-inflammatory not only in persisting viral infections (18), but also in other chronic diseases, such as various tumor types, where COX2 and/or PGE_2_ expression by tumors or myeloid derived suppressor cells have been correlated with suppression and deviation of anti-tumor T cell responses, as well as with worse disease outcome (19, 41). Currently there are great needs for drugs that can complement the existing checkpoint inhibitors in order to improve the proportion of patients showing clinical benefit. Interestingly, combined inhibition of PD-L1 and COX2 showed additive effects for the recovery of CD8+ T cell immunity in both models of chronic infection and tumors, with apparently limited side effects (18, 19) suggesting COX inhibitors may be used in combination therapy. More specific drugs targeting either PGES, PGE_2_ or EP2/4 could be of even greater benefit as they do not affect the other four COX-dependent prostanoid pathways (42). While inhibition of the COX or PGE_2_ pathway may act on cells residing within the site of chronic inflammation, including tumor cells and cancer associated fibroblasts (43) our study indicates that such inhibitors will also interfere with the anti-inflammatory capacity of FRC inside SLO, presumably within T zones, and may permit an improvement of the cancer-immunity cycle (44), namely a revitalization of previously exhausted T cell clones that can then seed again the sites of inflammation or cancer and maintain an effective immune response.

In summary, we propose that LN FRC act as a rheostat restraining T cell responses in at least two different ways, with weak responses being prevented by the omnipresent PGE_2_ and stronger responses being dampened by both PGE_2_ and the inducible NO. During chronic T cell responses, LN FRC may have a particularly critical role in restraining them via prostanoid release, presumably in an attempt to resolve the chronic inflammation that can have damaging effects on tissue function. These findings extend and strengthen the concept of suppressive LN stroma that can fine-tune T cell responses. We propose that FRC contribute directly and indirectly to the control of adaptive immune responses by hematopoietic cells, deciding either between immunity and tolerance, and regulating the extent of T cell clonal expansion, type of differentiation and longevity of the immune response.

## Experimental procedures

### Mice

C57BL/6J mice were purchased from Harlan Olac (Netherlands). *Nos2^−/−^* mice (45) and OT-1 mice (46) were as described. COX2^ΔCCL19Cre^ mice and ROSA26-EYFP^CCL19Cre^ were generated by intercrossing COX2^flox/flox^ (24) with CCL19Cre (47) and/or ROSA26-EYFP mice (48), with Cre- littermate mice used as controls. For experiments 6-week-old or older mice were used, all on a B6 background. All mouse strains used were bred and maintained in the SPF facility of the University of Lausanne. All mouse experiments were authorized by the Swiss Federal Veterinary Office (authorization numbers VD1612.3, VD1612.4 and VD3196).

### Adoptive cell transfers and immunizations

For adoptive T cell transfer 1x 10^5^ lymphocytes isolated from spleen and pLN of OT-1 mice (CD45.1+) were transferred i.v. into recipient mice (CD45.2+). The next day mice were immunized by s.c. injections in 6 sites in the flank with 5, 50 or 150µg Ovalbumin (OVA; Sigma) diluted in Montanide ISA 25 (25% in PBS; Seppic) containing 50µg Poly(I:C) (InvivoGen; # tlrl-pic). For studying memory responses 0.2-0.5 x 10^5^ OT-1 cells were transferred i.v. into recipient mice followed by a s.c. immunization at 2 sites with 50µg OVA/Montanide containing 10 or 25µg Poly(I:C), with the re-challenge performed d37 later with 5µg OVA/Montanide containing 5µg Poly(I:C), again s.c. into two sites.

### Viral infection

LCMV clone 13 virus stocks were generated according to an established protocol (49). To obtain a chronic infection 2 x10^6^ plaque forming units (PFU) of LCMV were injected i.v. and organs collected 8, 19 or 21 days post infection. To determine viral titers blood and organ samples were shock frozen on dry ice; tissues were homogenized by bead beating and viral titers determined by a focus-forming assay (49).

### Stromal and hematopoietic cell isolation

In order to investigate lymph node stromal cells, tissues were collected and digested as described elsewhere (3). Brief, peripheral LNs (axillary, brachial and inguinal) were removed and digested for 30min at 37°C under continuous stirring in DMEM medium containing 2% FCS, 3µg/ml Collagenase IV (Worthington) and DNAseI (Roche). Single cell suspensions of hematopoietic cells were obtained by meshing pLN and spleen through a 40µm nylon cell strainer. In spleen samples, erythrocytes were lysed with a Tris-Ammonium chloride based buffer.

### Cell lines

The wt FRC cell line pLN2 was described previously (11) and stems from naïve pLN of B6 mice. iNOS*^−/−^* FRC cell lines were generated accordingly from pLN of iNOS*^−/−^* mice. All cell lines were cultured in RPMI complete medium (RPMI 1640, 10%FCS, 10mM HEPES, 50µM β-Mercaptoethanol, P/S). For experiments cells were used between passages (p) 24- 32 (pLN2) and p23-27 (iNOS*^−/−^* FRC), respectively.

### *In vitro* T cell activation assay using agonistic antibodies

To investigate the effect of FRC on T cell activation co/cultures of cells were performed as described previously (11). Briefly, pLN2 or NOS2^−/−^ FRC lines were plated at 2.500 cells per 96 well and after overnight culture irradiated with 1000rad to prevent their rapid growth. T cells isolated from spleen and pLN (axillary, brachial and inguinal) of wt B6 mice were enriched by panning using antibodies to B220 (RA3-6B2), CD11b (M1/70) and CD11c (N418). These T cells were labeled with 2 µM CFSE (Invitrogen), resuspended in RPMI complete medium containing 3ng/ml murine IL-7 (Peprotech), 10U/ml human IL-2 (Merck Serono) and non-essential amino acids (Gibco) and plated at 2.5 x 10^5^ cells per 96-well, with or without an adherent layer of FRC. To activate T cells 1.25 x 10^5^ αCD3/CD28 coupled DynaBeads (Invitrogen) or 2.5 x 10^5^ MACSiBeads (Miltenyi Biotec) coupled with different amounts of αCD3/28 were added to the culture and T cell proliferation investigated on d3. Where indicated, inhibitors or the corresponding solvent control were added at d0 of co-culture; for the EP2 and EP4 antagonists a second bolus was added on d1. Inhibitor experiments used 10µM indomethacin (Sigma), 1µM 1400W dihydrochloride (Sigma), 5µM L161.982 (EP4 antagonist) and 5µM AH6809 (EP2 antagonist) (both Cayman Chemicals). Live cells were counted after 3d of co-culture and analyzed by flow cytometry. Percentage inhibition of T cell proliferation mediated by FRC was calculated based on the number of proliferated CD8+ and CD4+ T cells in the presence or absence of FRC in the co-culture.

### *In vitro* T cell activation using peptide-loaded BM-DC

As previously described (11), 1 x10^6^ BMDCs were activated with 0.5 µg/ml LPS (Sigma) for 6h at 37°C. 2h after LPS addition cells were loaded with titrated doses of N4 (SIINFEKL) or V4 (SIIVFEKL) peptide. Lymphocytes were isolated from spleen and pLN of wt B6 and OT-I mice, erythrocytes lysed, and CD8 T cells enriched by panning using antibodies to B220 (RA3-6B2), CD4 (H129.19.6), CD11b (M1/70) and CD11c (N418) followed by labeling with 2 µM CFSE (Invitrogen). Per 24-well 1.96×10^6^ wt T cells, 0.04×10^6^ CFSE^+^ OT-I T cells and 1×10^4^ BM-DC were cultured alone or in the presence of 1 x 10^4^ irradiated pLN2 FRC. After 3d of co-culture cells were harvested and analyzed by flow cytometry.

### Flow Cytometry

1-2 x 10^6^ cells were blocked with anti-CD16/32 antibody (2.4G2) and then stained for 40 min at 4°C using the following fluorochrome- coupled antibodies: CD44 (IM7), CD62L (MEL-14), IFNγ (XMG1.2), PD-1/CD279 (RMP1-30), TNFα (MP6-XT22), TCRβ (H57-597) were from eBioscience; CD45pan (30-F11), CD45R/B220 (RA3-6B2), CD8α (53-6.7), CD25 (PC61), CD31 (390) from Biolegend; CD19 (ID3), CD157 (BP-3) from BD Biosciences. CD4 (H129.19.6) and Podoplanin (8.1.1) antibodies have been generated in house. For intracellular staining, cells were fixed and permeabilized using the BD Cytofix/Cytoperm kit. Endogenous CD8^+^ T cells specific for LCMV were labeled with APC or PE conjugated peptide-MHC tetramers (Db/gp33-41, Db/gp276-286 and Db/NP396-404; TCMetrix) for 90min at 4°C. Dead cells were excluded by marking them with 7-AAD or the Fixable Aqua Dead Cell Stain Kit (both Invitrogen). Samples were acquired on a BD LSRII Flow Cytometer from BD Biosciences followed by analysis using FlowJo software (FlowJo LLC). Cell sorting was performed on a FACS-Aria (Becton Dickinson) using a 100µm nozzle. Different cell populations of lymph nodes were identified as previously described (3). For RNA isolation cells were directly sorted into lysis buffer (RNeasy micro kit, Qiagen).

### Re-stimulation and *in vitro* cytotoxicity assay

Lymphocytes from spleen or pLN of LCMV infected animals were isolated and 2 x10^6^ cells were seeded per 96-well. Cells were stimulated *in vitro* with 1µM of gp33 (KAVYNFATC) or np396 (FQPQNGQFI) peptide (ECM Microcollections) for 30 minutes at 37°C before BrefeldinA (10µg/ml, AppliChem) was added. After another 4h of incubation at 37° cells were harvested, washed and stained for surface or intracellular epitopes followed by analysis using flow cytometry. *In vitro* cytotoxicity assays were performed as previously described (11).

### Histology

Tissues of infected animals were fixed with 2% PFA for at least 4h, followed by overnight incubation in 30% sucrose before embedding in O.C.T. (Sakura Finetek). 8µm cryosections were generated and immunofluorescence staining performed as described recently (3). To monitor histological changes in pLN2 or iNOS^−/−^ FRC lines in 8-well chamber slides (Falcon) 7.5 x 10^3^ irradiated FRC were seeded per chamber (Falcon), and then superseeded by 3.75 x 10^5^ T lymphocytes harvested from spleen and pLN of wt mice and activated with 3.75 x 10^5^ αCD3/28 coated MACSiBeads. After 2d of co-culture T cells were removed and adherent FRCs stained for microscopic analysis. Cells were fixed with 100% cold acetone and staining was performed as described above. Images were acquired with a Zeiss Axioplan microscope and treated with Photoshop (Adobe) or ImageJ software respectively.

The following antibodies were used for histology: CD3 (145-2c11), CD31 (GC-51), CD35 (8C12), B220 (RA3-6B2), CD157 (BP 3.4), Podoplanin (8.1.1) antibodies have been generated in house;

CCL21 (AF457, R&D Systems), Lyve-1 (103-PA50, RELIATech), Laminin (L9393, Sigma), iNOS (06-573, Millipore), IgG+IgM (315-005-048, Jackson).

### Quantitative Realtime PCR

To investigate the genetic profile of stroma versus hematopoietic cell enriched tissue fractions, spleen or pLN were meshed through a 40µm filter (4). The non-soluble stroma remaining in the filter was harvested directly in TRIzol (Ambion, life technologies) whereas the soluble hematopoietic cells were centrifuged and then resuspended in TRIzol. These Trizol samples were homogenized by bead beating and RNA extracted (3). RNA of sorted cells was isolated using RNeasy Micro Kit (Qiagen). Reverse transcription and quantitative real-time PCR was performed as described previously (3). Efficiency-corrected RNA expression was normalized to the expression of the two housekeeping genes *hprt* and *tbp*. Sequences of primer pairs used are as follows:

*inos* (NOS2): Fwd: 5’- GTT CTC AGC CCA ACA ATA CAA GA −3’ and Rev: 5’- GTG GAC GGG TCG ATG TCA C-3’

*ptgs1* (COX1): Fwd: 5’- CCA GAG TCA TGA GTC GAA GGA G −3’ and Rev: 5’- CCT GGT TCT GGC ACG GAT AG −3’

*ptgs2* (COX2): Fwd: 5’-AAT TAC TGC TGA AGC CCA CC-3’ and Rev: 5’-CTT CCC AGC TTT TGT AAC CAT-3’

*ptges1*: Fwd: 5’-GCA CAC TGC TGG TCA TCA AG-3’and Rev: 5’- ACG TTT CAG CGC ATC CTC-3’

*ptges2*: Fwd: 5’- CAG GTG GTG GAG GTG AAT CC-3’ and Rev: 5’- CTG CCC TGA AAC CAG GTA GG-3’

*ptges3*: Fwd: 5’- CGA ATT TTG ACC GTT TCT CTG-3’ and Rev: 5’- TGA ATC ATC ATC TGC TCC ATC T-3’

*hprt*: Fwd: 5’- GTTGGATATGCCCTTGAC-3’ and Rev: 5’-AGG ACT AGA ACA CCT GCT-3’

*tbp*: Fwd: 5’- CCT TCA CCA ATG ACT CCT ATG AC-3’ and Rev: 5’- CAA GTT TAC AGC CAA GAT TCA C-3’

### Prostaglandin E_2_ and Nitrite detection

PGE_2_ levels were determined in culture supernatants using the PGE_2_ ELISA Kit (Cayman Chemical) according to the manufacturer’s instructions. The level of NO_2_ in cell culture supernatants was measured using the Griess Assay (11).

### Statistical analysis

Statistical analyses were performed with Prism (version 6.0b and 7.0d, GraphPad Software). Unpaired t-test (normally distributed data) or Mann-Whitney test (for not normally distributed data) were used to compare two data sets. To compare multiple groups, One-Way Kruskal-Wallis non-parametric tests were performed followed by a post-hoc test (Dunn’s test). P values < 0.05 were considered to be statistically significant.

## Author contributions

Conceived and designed experiments: SAL, KS, DZ. Performed experiments: KS, HC, SF, HYH, SGO. Analyzed data: KS, SAL, HC, SF. Contributed reagents or virus stocks: SAL, DZ. Wrote the manuscript: SAL, KS, with critical input by all authors.

## Acknowledgments

We thank: Stefanie Siegert, Leonardo Scarpellino, Francois Renevey, Patricia Aparicio, Tobias Vogt, Loriane Savary, Mégane Bernard, Francesca Alfei, Stefanie Scherer, Florence Morgenthaler (Cellular Imaging Facility), Anne Wilson and Danny Labes (Flow cytometry facility) for advice and technical assistance; Daniel F. Legler for reagents and advice; Harvey Hershman (UCLA) and Gerhard Püschel for providing the COX2^flox/flox^ mice and Burkhard Ludewig (Kantonsspital St.Gallen) for CCL19cre mice; Daniel Speiser for critical reading of the manuscript. This study was supported by grants from the Swiss National Science Foundation (31003A-166161 to SAL), the Novartis Foundation (to KS and SAL) and from the European Research Council (ProtecTC to DZ). The authors have no conflicting financial interests.

## Figure Legend for Supplementary data

**Supplementary Figure 1:**
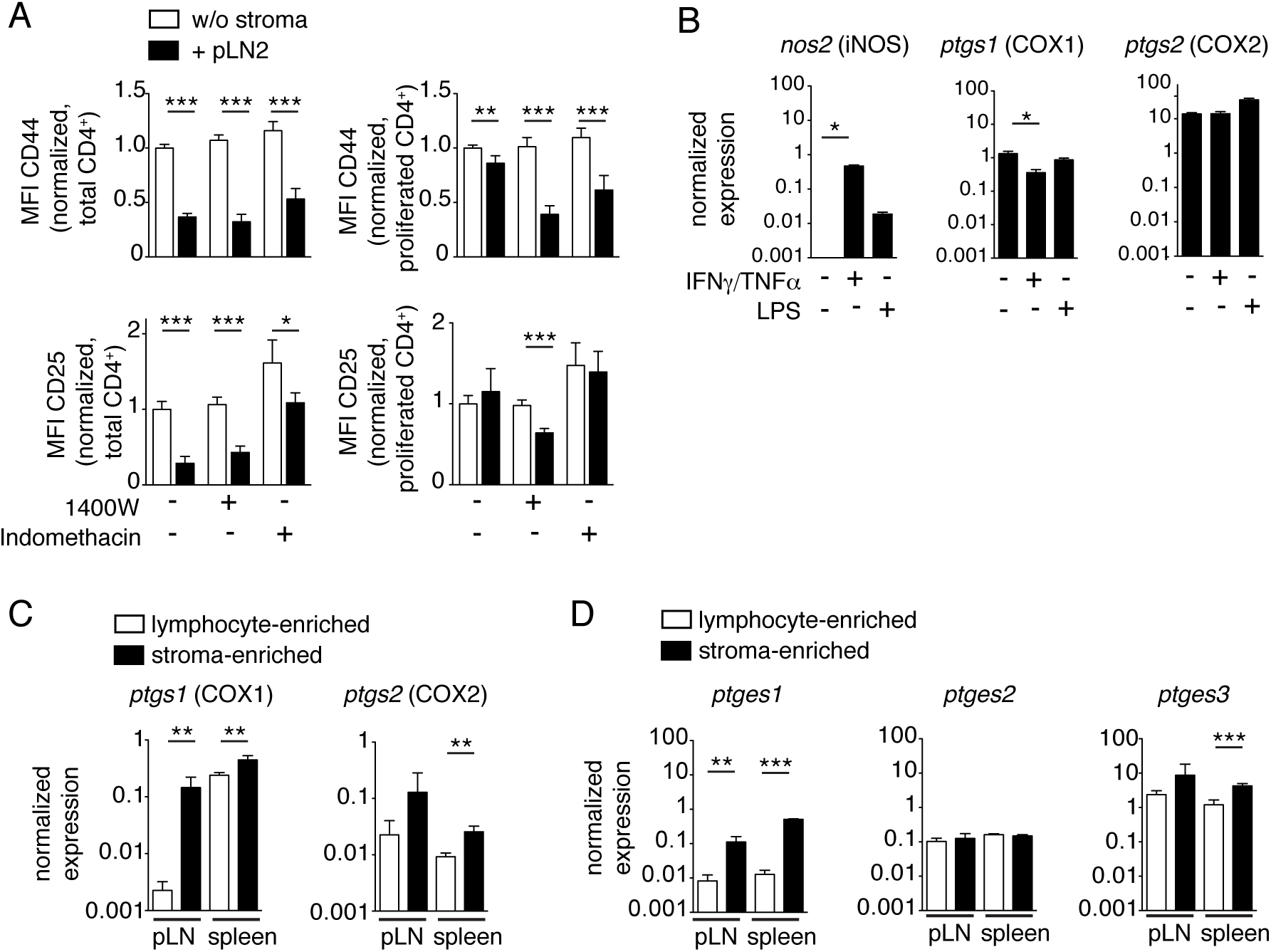
Activation of CD4+ T cells is dampened by iNOS and COX2 activity in pLN FRC. (A) T cells were activated with αCD3/28 DynaBeads and cultured for 3d in the absence (white bars) or presence of pLN2 (black bars) ± 1400W (3µM) ± indomethacin (10µM). Flow cytometric analysis showing the MFI of CD25 and CD44 on total or proliferated CD4+ T cells (pool of two independent experiments, n=4). Bar graphs showing mean ± STD. (B) qRT-PCR analysis for *Nos2*, *ptgs1* and *ptgs2* transcript levels in pLN2 cells that were left unstimulated or stimulated for 7h with 10ng/ml IFNγ/TNFα or 0.5µg/ml LPS (n=3). (C-D) qRT-PCR analysis of the soluble (lymphocyte enriched) and non-soluble (stroma enriched) fractions of pLN and spleen of naïve wt mice (n=4) for transcripts of *ptgs1/2* (C) or *ptges1/2/3* (D). Bar graphs show the mean ± STD. Statistics in (A, C and D) using unpaired t-test and in (B) using a Kruskal-Wallis followed by Dunn’s multiple comparison test. \**P <* 0.05, \*\**P <* 0.005 and \*\*\**P <* 0.001

**Supplementary Figure 2:**
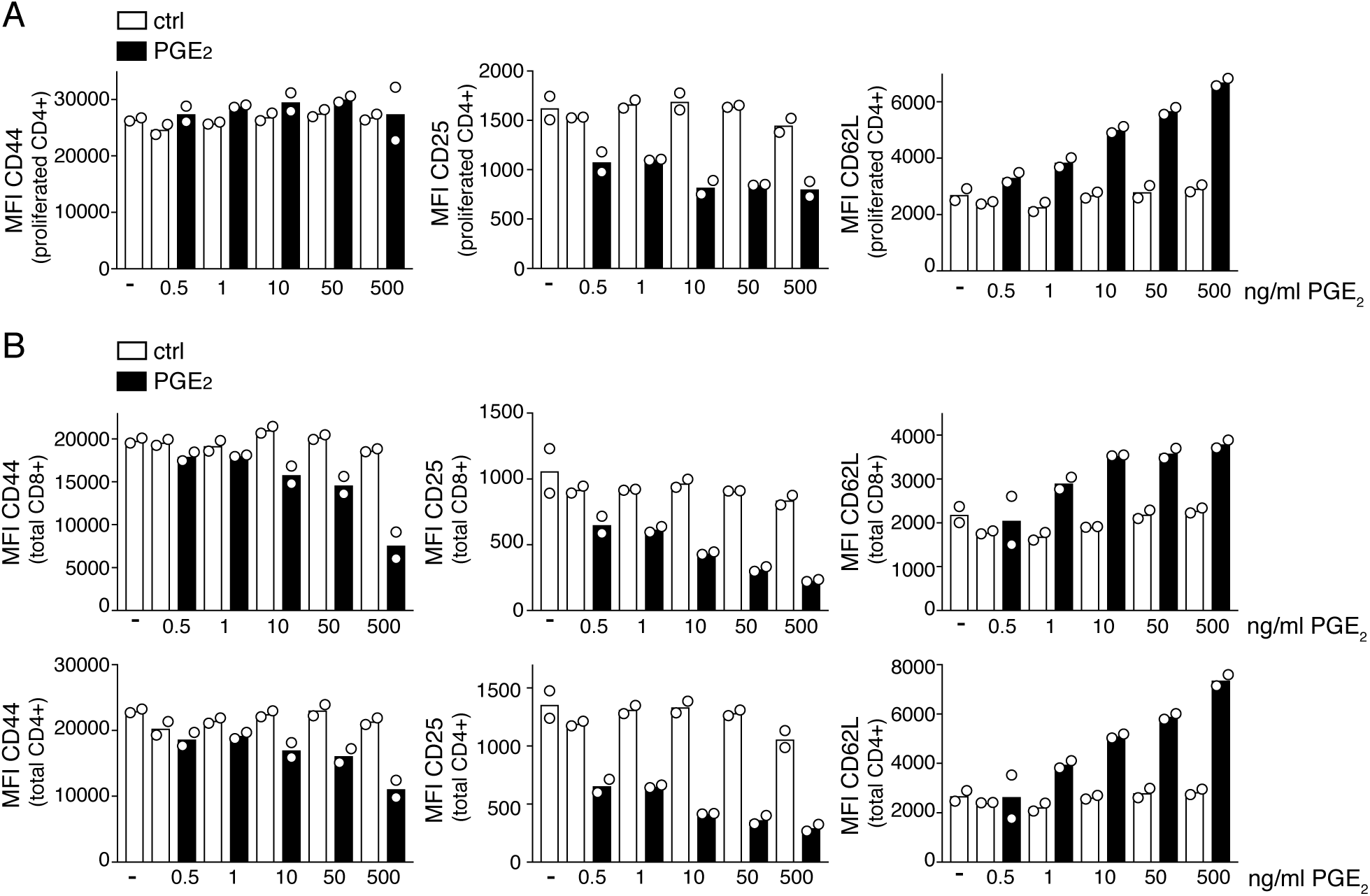
PGE_2_ dampens effector cell phenotype in activated T cells. Flow cytometric analysis of T cells activated with αCD3/28-coated beads and cultured for 3 days with the indicated concentrations of PGE_2_ (black bars) or solvent control (white bars), respectively. MFI of CD44, CD25 and CD62L on proliferated CD4+ T cells (A) and total CD8+ or CD4+ T cells (B). Proliferation was assessed by CFSE dilution. Data are representative of 3 independent experiments. Bar graphs show mean ± STD.

**Supplementary Figure 3:**
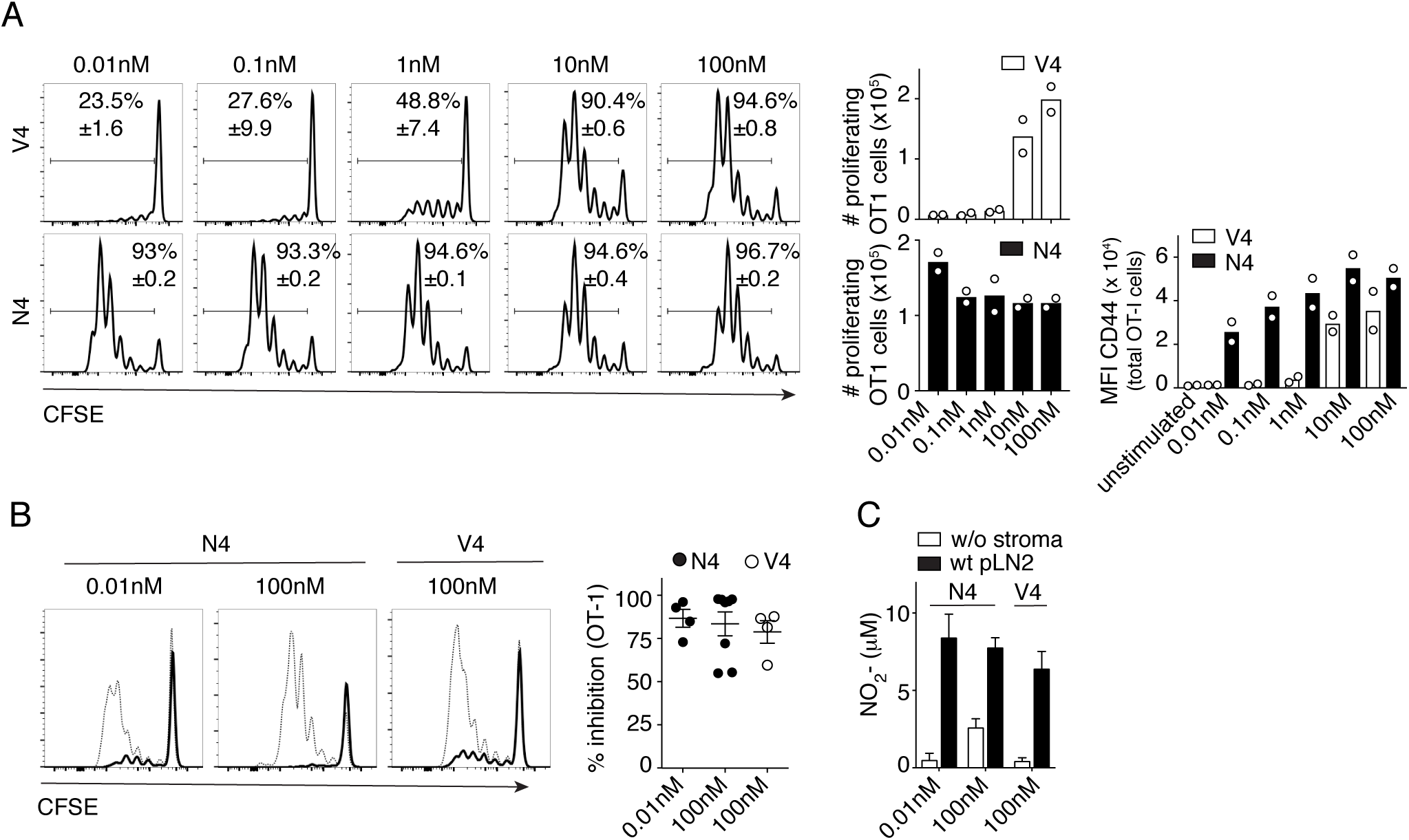
Lymph node FRC dampen the expansion of CD8+ T cells of high and low antigen affinity. CFSE-labeled OT-1 CD8+ T cells were mixed in a ratio of 1: 50 with wt T cells and cultured with LPS-activated BM-DC pulsed with the indicated concentrations of OVA peptides of high affinity (N4) or low affinity (V4) for the OT-1 receptor. Co-cultures were performed in the absence or presence of wt pLN2 FRC and OT-1 cell proliferation (A, B) or nitrite levels (C) were assessed after 3d of culture. (A) CFSE profiles (left side), numbers (middle panel) and CD44 expression levels (right panel) of OT-1 T cells activated in the absence of the pLN2 FRC line. Data are representative of 2 independent experiments performed in duplicates. (B) CFSE profile of OT-1 cells cultured in the absence (thin line) or presence (black line) of pLN2 FRC. Scatter dot plot depicts the percentage inhibition of OT-1 T cell proliferation by FRC. (C) Bar graphs showing nitrite (NO2-) level found in the supernatant of the co-cultures shown in (B). Data in B and C represent a pool of two independent experiments; n≥4.

**Supplementary Figure 4:**
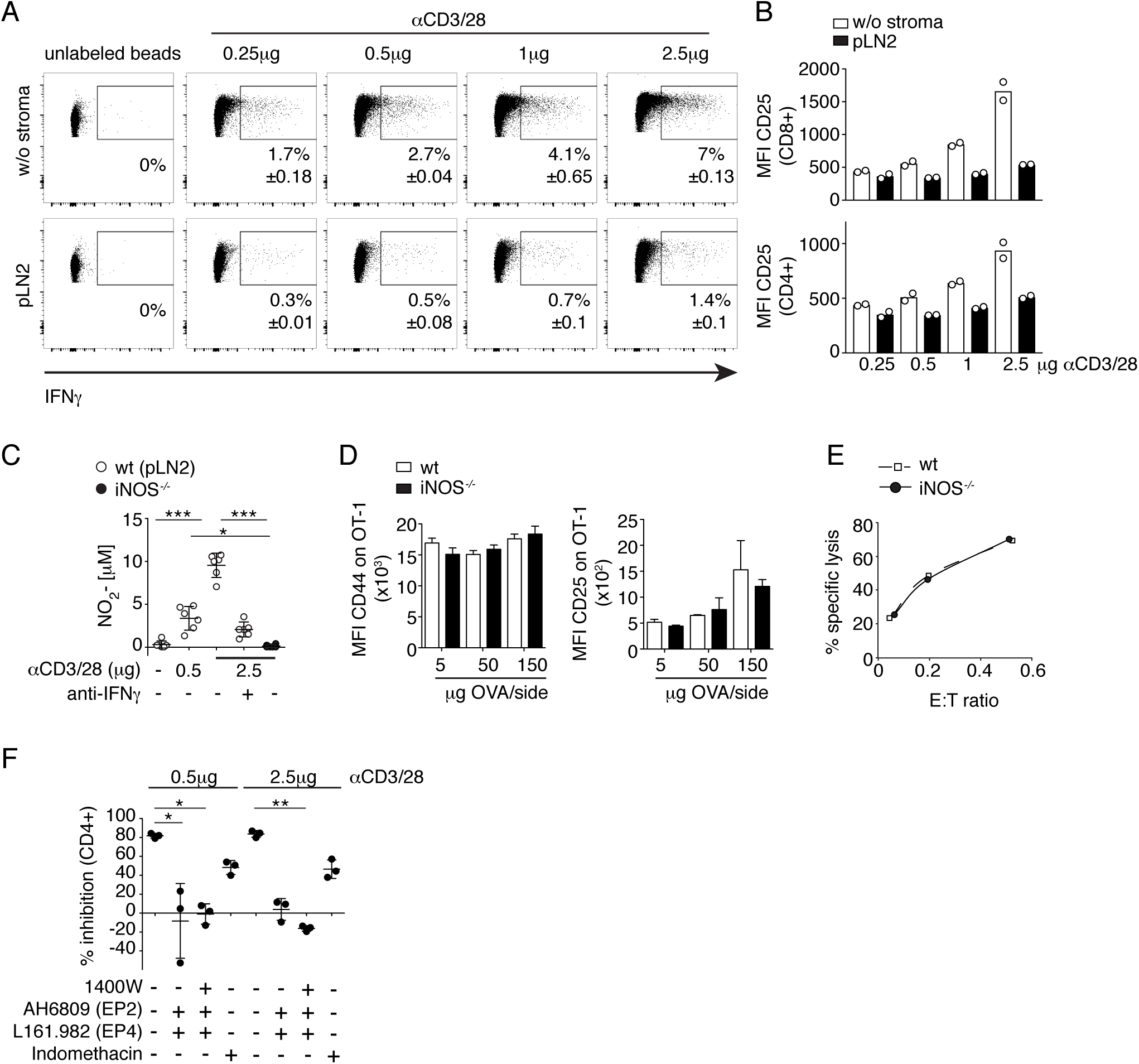
The magnitude of iNOS-mediated T cell inhibition correlates with increased T cell strength and early IFNγ production but does not impact effector function of proliferating cells. (A-C, F) CD8+ and CD4+ T cells were activated with the indicated amount of αCD3/28 coated on MicroBeads, in the absence or presence of pLN2 cells. Frequency of IFNγ-producing CD8+ T cells (A) and expression levels of CD25 on CD8+ and CD4+ T cells (B) were investigated after 1d of co-culture. 1 representative out of 3 independent experiments is shown, with at least two replicates each. (C) iNOS*^−/−^* and wt (pLN2) FRC cell lines were co-cultured with activated T cells ± neutralizing anti-IFNγ antibodies and nitrite levels measured in the SN of d2 cultures using the Griess assay. Scatter plot showing 1 representative out of 3 independent experiments. (D) Wt or iNOS*^−/−^* mice received OT-1 CD8+ T cells i.v. and were then immunized s.c. with the indicated concentrations of OVA/Montanide with draining pLNs investigated on d4 after immunization. Bar graphs depict the MFI of CD44 and CD25 expression on OT-1 T cells isolated from wt or iNOS*^−/−^* mice (n≥8, pool of 2-3 independent experiments). (E) Cytotoxic capacity of OT-1 T cells isolated from draining pLN of wt and iNOS*^−/−^* mice immunized s.c. 4 days earlier with 150ug OVA/Montanide. Shown is the percentage of target cell lysis with the indicated Effector:Target (E:T) ratios for one representative (n=3) out of 2 independent experiments. (F) Cocultures of T cells activated with beads coated with either 0.5 or 2.5µg CD3/28 antibody and cultured for 3d in the presence or absence of pLN2 FRC ± indicated inhibitors (1400W (3µM), AH6809 (5µM); L161.982 (5µM); indomethacin (10µM). Scatter plot depicts the percentage inhibition of CD4+ T cell expansion mediated by FRC (n=3; data representative of 3 independent experiments). Statistics: (C and F) Kruskal Wallis followed by Dunns post-test; \**P <* 0.05, \*\**P <* 0.005 and \*\*\**P <* 0.001

**Supplementary Figure 5:**
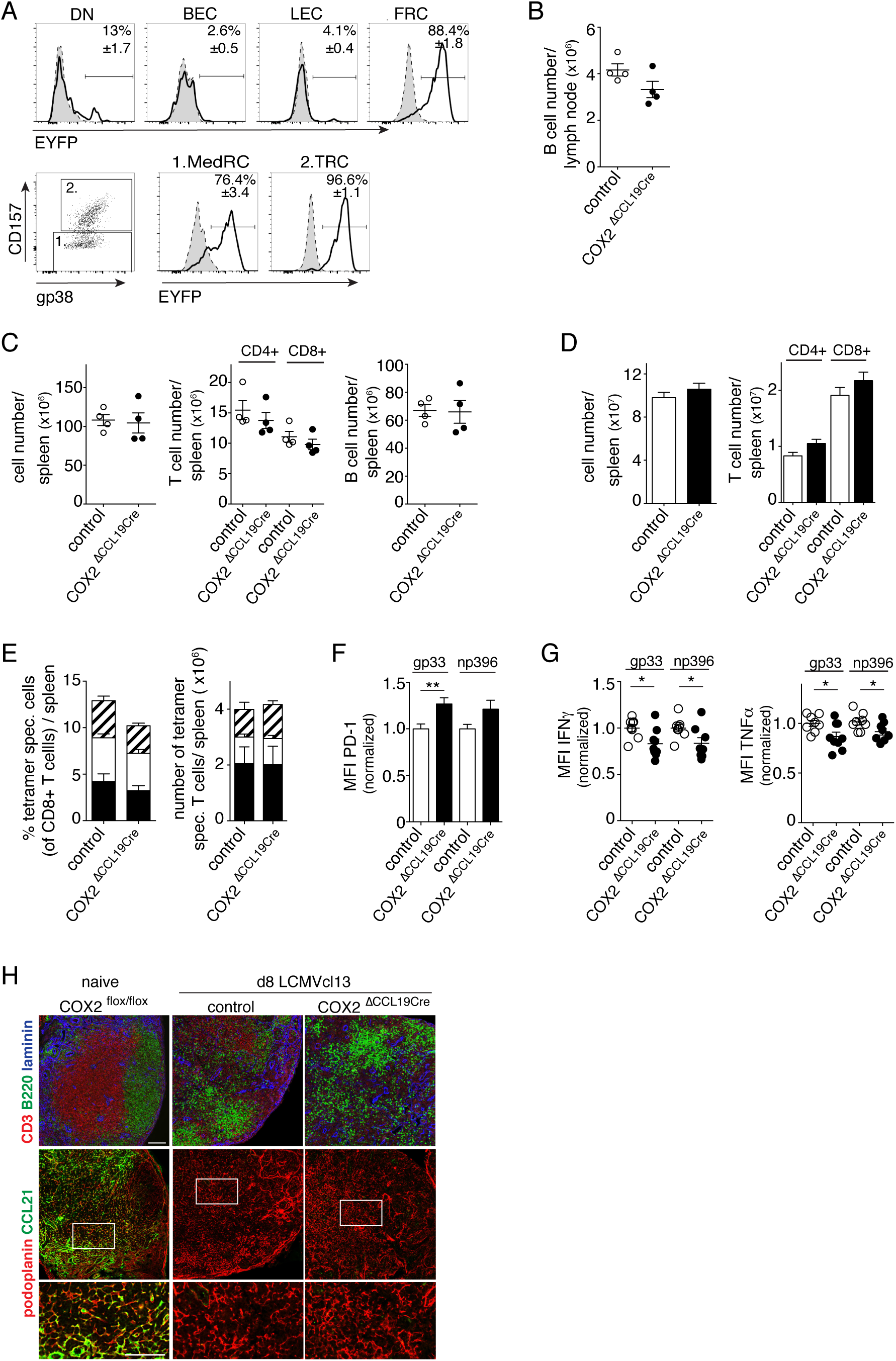
COX2 deletion in FRC has no influence on the cellular composition and organization of pLN and spleen. Flow cytometric (A-F) and histological (G) analysis of COX2^ΔCCL19Cre^ mice and Cre- littermates, either naive (A-C,H) or 8d after infection with LCMV clone 13 (D-H). (A) CCL19-Cre activity was investigated in pLN of naïve COX2^ΔCCL19Cre^ ROSA26-EYFP^CCL19Cre^ mice. Histograms showing EYFP expression in different non-hematopoietic cell types (CD45-): CD31- podoplanin- double-negative (DN) cells, CD31+ podoplanin- blood endothelial cells (BEC), CD31+ podoplanin+ lymphatic endothelial cells (LEC) and CD31- podoplanin+ FRC in EYFP-reporter (solid black line) compared to control Cre- mice (gray shading). FRCs were further divided into CD157- Medullary FRC (MedRC) and CD157+ T-zone FRC (TRC) (n=3, representative data of 2 independent experiments). (B-C) Scatter plots showing CD19+ B cell numbers in pLN (B) and lymphocyte numbers in the spleen of COX2^ΔCCL19Cre^ (black circles) and Cre- littermate controls (white circles; ‘control’). Data in (B) and (C) show 1 representative out of 3 experiments. (D-H) COX2^ΔCCL19Cre^ and Cre- littermate mice were infected with 2×10^6^ PFU LCMV clone 13 and the spleen analyzed on d8 post infection. (D) Bar graphs showing total cell numbers and T cell numbers as well as (E) frequencies and cell numbers of LCMV-specific CD8+ T cells (specific for three viral peptides) (pool of two independent experiments; n≥7). (F) Median fluorescent intensity (MFI) of PD-1 on LCMV-specific CD8+ T cells, with cells from Cre+ mice normalized to those from Cre- mice. (G) Frequencies of IFNγ– or TNFα– expressing LCMV-specific cells determined after short re-stimulation with gp33 or np396 peptides, respectively. Scatter plot showing normalized frequencies of LCMV-specific cells of Cre+ compared to Cre- controls (Data in F and G show a pool of two independent experiments; n≥5). (H) Immunofluorescence microscopy analysis of labeled pLN sections from naïve or d8 LCMV clone 13 infected mice of the indicated genotype. Representative images from 3 mice/genotype are depicted. Scale bar, 100µm. Bar graphs and scatter plots showing mean ± SEM. Statistics: (F-G) unpaired t-test; \**P <* 0.05, \*\**P <* 0.005 and \*\*\**P <* 0.001

**Supplementary Figure 6:**
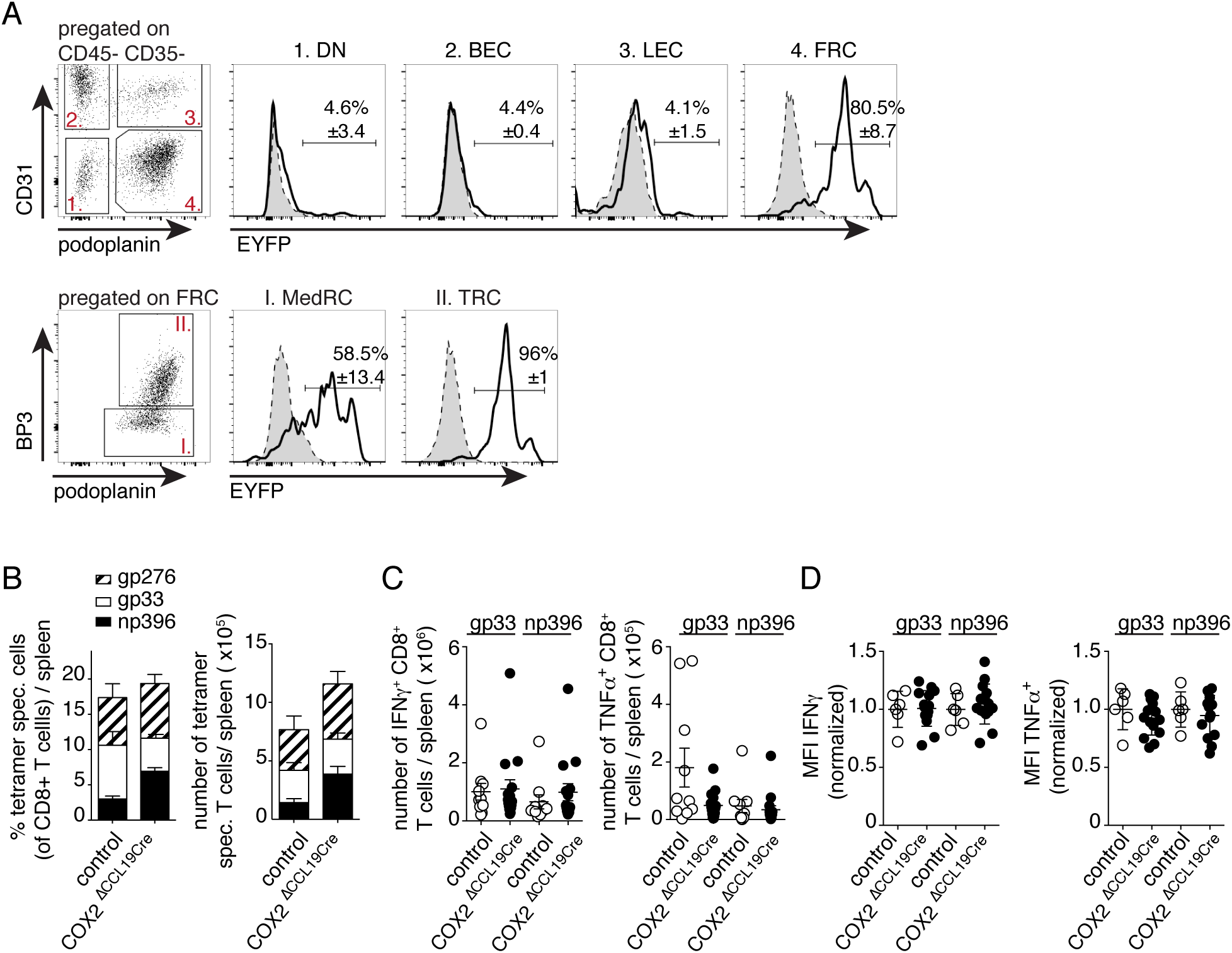
COX2 depletion in FRC leads to an increased virus-specific CD8+ T cell response during the chronic phase of LCMV clone 13 infection. (A) COX2^ΔCCL19Cre^ ROSA26-EYFP^CCL19Cre^ and Cre- littermate mice were infected with LCMV clone 13 and digested pLN analyzed on d19 post infection. Cre activity was investigated by measuring EYFP levels in different cell types using flow cytometry. Dot plot shows gating strategy to distinguish CD31- podoplanin- DN cells, CD31+ podoplanin- BEC, CD31+ podoplanin+ LEC and CD31- podoplanin+ FRC after pregating on CD45- CD35- non-hematopoietic cells. The FRC population was further subdivided into CD157- MedRC and CD157+ TRC. The histograms show the frequency of EYFP expressing cells among different stromal cell subsets, with the cells from Cre- mice shown with gray shading. 1 out of two representative experiments is shown (n=3). (B-D) The spleens of COX2^ΔCCL19Cre^ and control mice were analyzed on d21 post clone 13 infection for the frequency and number of LCMV-specific CD8+ T cells (B), for the frequency of IFNγ- or TNFα–expressing splenic CD8+ T cells (C), or for the MFI of intracellular IFNγ or TNFα levels in CD8+ T cells (D; normalized to controls), after re-stimulation with gp33 or np396 peptides, respectively (C, D: pool of three independent experiments; n≥6). (B-D) Bar graphs and scatter plots showing mean ± SEM.

